# Methyl-CpG-binding domain 9 (MBD9) is required for H2A.Z incorporation into chromatin at a subset of H2A.Z-enriched regions in the Arabidopsis genome

**DOI:** 10.1101/404152

**Authors:** Paja Sijacic, Dylan H. Holder, Marko Bajic, Roger B. Deal

**Author notes:** Correspondence: Roger B. Deal.

## Abstract

The SWR1 chromatin remodeling complex, which deposits the histone variant H2A.Z into nucleosomes, has been characterized in yeast and animals but had not been purified from plants. We used the conserved SWR1 subunit ACTIN RELATED PROTEIN 6 (ARP6) as bait in tandem affinity purification experiments to isolate associated proteins from *Arabidopsis thaliana*. We identified all 11 subunits found in yeast SWR1 and the homologous mammalian SRCAP complexes, demonstrating that this complex is conserved in plants. We also identified several additional proteins not previously associated with SWR1, including Methyl-CpG-BINDING DOMAIN 9 (MBD9). Since *mbd9* mutant plants were phenotypically similar to *arp6* mutants, we further explored a potential role for MBD9 in H2A.Z deposition. We found that MBD9 is required for proper H2A.Z incorporation at thousands of discrete sites, which represent a subset of the regions normally enriched with H2A.Z. Genetic analyses showed that *arp6;mbd9* double mutants have far more severe phenotypes than either single mutant. In conjunction with the finding that MBD9 does not appear to be a core subunit of the Arabidopsis SWR1 complex, this suggests that MBD9 also has important roles beyond H2A.Z deposition. Our data establish the SWR1 complex as being conserved across eukaryotes and also provide new insights into the mechanisms that target H2A.Z to chromatin.

## INTRODUCTION

Nucleosomes, the fundamental repetitive units of chromatin that consist of ∼147 bp of DNA wrapped around a histone octamer, efficiently condense large eukaryotic DNA molecules inside the nucleus. At the same time, nucleosomes represent a physical barrier that restricts the access of DNA-binding proteins to regulatory sequences. This physical constraint imposed by nucleosomes on DNA can be modulated to expose or occlude regulatory DNA sequences, and is thereby used as a mechanism to control processes such as transcription that rely on sequence-specific DNA binding proteins. Thus, enzymatic complexes that can remodel chromatin structure by manipulating the position and/or composition of nucleosomes are essential for proper transcriptional regulation and the execution of key developmental programs.

All chromatin-remodeling complexes (CRCs) contain a DNA-dependent ATPase catalytic subunit that belongs to the SNF2 family of DNA helicases, along with one or more associated subunits (Adam et al., 2001; Hota and Bruneau, 2016). There are four major subfamilies of CRCs: SWI/SNF, INO80, ISWI, and CHD, all of which use the energy of ATP to either slide, evict, or displace nucleosomes, or to replace the canonical histones within nucleosomes with histone variants. One member of the INO80 CRC subfamily is the SWI2/SNF2-related 1 (SWR1) chromatin remodeler, a multisubunit protein complex required for incorporation of the H2A variant, H2A.Z, into chromatin (Kobor et al., 2004; Mizuguchi et al., 2004). H2A.Z is a highly conserved histone variant found in all eukaryotes that plays important roles in regulating a variety of cellular processes, including transcriptional activation and repression, maintenance of genome stability and DNA repair, telomere silencing, and prevention of heterochromatin spreading (Jarillo and Pineiro, 2015; March-Diaz and Reyes, 2009; Marques et al., 2010; Meneghini et al., 2003; Raisner and Madhani, 2006; Rosa et al., 2013; Xu et al., 2012; Zhou et al., 2010). Although mutations in H2A.Z are not lethal in yeast (Adam et al., 2001; Jackson and Gorovsky, 2000), H2A.Z is essential for viability in other organisms such as Tetrahymena (Liu et al., 1996), Drosophila (Clarkson et al., 1999; van Daal and Elgin, 1992), and mice (Faast et al., 2001). Interestingly, H2A.Z-deficient plants are viable but display many developmental abnormalities such as early flowering, reduced plant size, altered leaf morphology, and reduced fertility (Choi et al., 2007; Coleman-Derr and Zilberman, 2012; March-Diaz et al., 2008).

The SWR1 complex that mediates H2A.Z incorporation into chromatin was first described in yeast and is composed of 13 subunits, including Swr1, the catalytic and scaffolding subunit of the complex (Kobor et al., 2004; Krogan et al., 2003; Mizuguchi et al., 2004; Wu et al., 2009). In mammals, the functional and structural homolog of yeast SWR1 complex is the SRCAP (SNF2-related CREB-binding protein activator protein) complex. This complex is composed of 11 of the same subunits found in yeast SWR1 and is likewise able to exchange H2A/H2B dimers for H2A.Z/H2B dimers in nucleosomes (Cai et al., 2005; Cai et al., 2006; Ruhl et al., 2006; Wong et al., 2007). Intriguingly, higher eukaryotes possess an additional multisubunit complex, called dTIP60 in Drosophila and TIP60 in mammals, that has histone acetyltransferase (HAT) activity and can also mediate the deposition of H2A.Z into nucleosomes (Doyon et al., 2004; Gevry et al., 2007; Kusch et al., 2004; Martinato et al., 2008). Several components of the TIP60 complex are homologous to SWR1 subunits, including the largest subunit, p400/Domino, which is an evolutionary derivative of the yeast Swr1 ATPase and the mammalian SRCAP ATPase (Cai et al., 2005; Cai et al., 2003; Kusch et al., 2004; Lu et al., 2009). However, other subunits in the TIP60 complex are homologous to those of the yeast HAT complex called NuA4. In fact, the TIP60 complex was originally identified as the functional equivalent of the yeast NuA4, with a primary function of acetylating histones (Doyon et al., 2004; Kimura and Horikoshi, 1998; Kusch et al., 2004). In yeast, the SWR1 and NuA4 complexes cooperatively regulate the incorporation of H2A.Z into chromatin and are structurally related through the sharing of four subunits: SWR1 COMPLEX SUBUNIT 4 (SWC4), YEAST ALL1-FUSED GENE FROM CHROMOSOME 9 (YAF9), ACTIN-RELATED PROTEIN 4 (ARP4), and ACTIN1 (ACT1) (Altaf et al., 2010; Babiarz et al., 2006; Keogh et al., 2006; Lu et al., 2009; Millar et al., 2006).

It appears that in higher eukaryotes the TIP60 complex is a structural and functional equivalent of a merger of the SWR1 and NuA4 complexes. During evolution, higher eukaryotes combined these two complexes into a single multifunctional complex, whose primary function is to acetylate histone tails, but is also capable of depositing H2A.Z into nucleosomes (Lu et al., 2009). While mammalian SRCAP and TIP60 share the same four subunits as yeast SWR1 and NuA4: SWC4, YAF9, ARP4, and ACT1, some subunits are unique to one complex or the other. For instance, SRCAP, ARP6, and ZnF-HIT1 are subunits found exclusively in the SRCAP complex (Lu et al., 2009).

Many homologs of yeast SWR1 and animal SRCAP complex subunits have been identified in *Arabidopsis thaliana* including ACTIN RELATED PROTEIN 6 (ARP6), SWR1 COMPLEX SUBUNIT 2 (SWC2), SWC6, PHOTPERIOD-INDEPENDENT EARLY FLOWERING 1 (PIE1), and three H2A.Z paralogs: HTA8, HTA9, and HTA11. For instance, PIE1 was first identified as a homolog of the Swr1, SRCAP, Domino, and p400 ATPases since it contains the same functional domains found in these proteins (March-Diaz and Reyes, 2009). Numerous genetic and biochemical experiments suggest that the SWR1 complex is conserved in Arabidopsis. For example, it has been recently shown that Arabidopsis SWC4 protein directly interacts with SWC6 and YAF9a, two known components of the SWR1 complex (Gomez-Zambrano et al., 2018). Additionally, protein interaction experiments have demonstrated that PIE1 interacts directly with ARP6, SWC6, and three H2A.Z proteins: HTA8, HTA9, and HTA11 (Choi et al., 2007; Lazaro et al., 2008; March-Diaz et al., 2007; March-Diaz et al., 2008), suggesting that PIE1 serves as the catalytic and scaffolding subunit of an Arabidopsis SWR1-like complex. Furthermore, functional characterizations of *PIE1, ARP6* and *SWC6* have revealed that the mutations in these genes have similar pleiotropic effects on Arabidopsis development, including a loss of apical dominance, curly and serrated rosette leaves, early flowering due to reduced expression of *FLOWERING LOCUS C* (*FLC*), altered petal number, and reduced fertility (Choi et al., 2005; Choi et al., 2007; Deal et al., 2005; Deal et al., 2007; Lazaro et al., 2008; March-Diaz et al., 2007; Martin-Trillo et al., 2006; Noh and Amasino, 2003). Interestingly, genetic experiments revealed that the *pie1* null phenotypes are more severe than those of *arp6, swc6, hta9;hta11*, or *hta8;hta9;hta11* (*h2a.z* near-null) plants (Choi et al., 2005; Choi et al., 2007; Coleman-Derr and Zilberman, 2012; Deal et al., 2005; Deal et al., 2007; Lazaro et al., 2008; March-Diaz et al., 2007; Martin-Trillo et al., 2006; Noh and Amasino, 2003). The more dramatic phenotypes in *pie1* plants suggest that PIE1 has additional functions outside of H2A.Z deposition by SWR1, as previously proposed (Choi et al., 2007; Coleman-Derr and Zilberman, 2012; Jarillo and Pineiro, 2015; March-Diaz and Reyes, 2009). A recent report also showed that mutant plants null for *pie1* and *h2az* exhibited early developmental arrest, dying shortly after germination (Coleman-Derr and Zilberman, 2012), further supporting the notion that PIE1 has H2A.Z-independent functions in Arabidopsis. On the other hand, genetic analyses of *pie1;swc6* double mutant plants revealed that they had a phenotype indistinguishable from *pie1* single mutants (March-Diaz et al., 2007), and *arp6;swc6* plants displayed the same defects as either *arp6* or *swc6* single mutant plants (Choi et al., 2007; Lazaro et al., 2008). These results further support the idea that PIE1, ARP6, and SWC6 act in the same genetic pathway and/or are the components of the same protein complex, but that PIE1 has additional functions.

Despite the strong genetic and biochemical evidence that Arabidopsis contains many conserved subunits homologous to the components of the yeast SWR1 complex and mammalian SRCAP, the plant SWR1 complex has not been successfully isolated and characterized. Recently, Bieluszewski and colleagues used Arabidopsis SWC4 and ARP4 proteins, two shared subunits of yeast SWR1 and NuA4 and mammalian SRCAP and TIP60 complexes, as baits to affinity-purify their interacting partners from Arabidopsis cell suspension cultures (Bieluszewski et al., 2015). These studies identified most of the subunits normally found in the SWR1 and NuA4 complexes, but it was not possible to determine whether this collection of proteins represented a single large complex or multiple complexes. Bieluszewski and colleagues originally hypothesized that plants may contain a TIP60-like complex with PIE1 acting as a catalytic and scaffolding subunit, similar to p400 in mammalian TIP60. However, they also identified an Arabidopsis EAF1-like protein, a homolog of the yeast Eaf1 that serves as a platform subunit of the yeast NuA4 complex (Auger et al., 2008; Bieluszewski et al., 2015), suggesting that plants may also contain a separate NuA4-like complex. Overall, it is not yet clear whether plants possess separate SWR1 and NuA4 complexes, SWR1 and TIP60-like complexes assembled around PIE1, or all three complexes (Bieluszewski et al., 2015).

The main goal of our study was to purify the Arabidopsis SWR1 complex and to identify all of its components. To achieve this, we used the ARP6 protein, a subunit unique to SWR1 in other organisms, as bait in Tandem Affinity Purification (TAP) experiments. We performed three independent TAP experiments to isolate and identify ARP6-associated proteins. We identified all 11 conserved subunits found in yeast SWR1 and mammalian SRCAP complexes, demonstrating that Arabidopsis contains a bona-fide functional and structural homolog of these complexes. In addition, we identified several unexpected proteins that associated with ARP6, including the plant homeodomain (PHD)- and Bromo domain-containing protein Methyl CpG-BINDING DOMAIN 9 (MBD9), and the PHD domain-containing proteins ALFIN-LIKE 5 (AL5), AL6, and AL7. Genetic analyses revealed that *mbd9* mutants showed phenotypic similarities to *arp6* mutants, so we further explored a possible role for MBD9 in regulating H2A.Z incorporation into chromatin. We found that MBD9 is required for H2A.Z incorporation at a subset of the sites that normally harbor H2A.Z nucleosomes, and that these MBD9- dependent H2A.Z sites have distinct chromatin features. Further, we found that MBD9 is not a core subunit of the Arabidopsis SWR1 complex and that double mutant *arp6;mbd9* plants exhibited much more severe phenotypes than single *arp6* or *mbd9* mutants. These results collectively suggest that MBD9 targets the SWR1 complex to a subset of genomic loci but also has important functions beyond H2A.Z deposition.

## RESULTS

### ARP6 transgenes tagged with a Tandem Affinity Purification tag rescue the *arp6-1* phenotype

To isolate the Arabidopsis SWR1 complex, we decided to use ARP6 protein as bait in Tandem Affinity Purification (TAP) experiments because ARP6 is exclusively found in the SWR1 complex in other organisms and is not shared by any other known CRCs that regulate H2A.Z incorporation into chromatin (Choi et al., 2007; Lu et al., 2009; Mizuguchi et al., 2004). For our purification experiments we used a GS^rhino^ TAP-tag, which consists of two protein G domains, a tandem repeat of rhinovirus 3C protease cleavage site, and the streptavidin-binding peptide. This tag has been successfully used to purify several plant nuclear complexes, including SWI/SNF type CRCs (Van Leene et al., 2015). Furthermore, the use of this tag provides high yield of purified proteins and specificity of purification. In addition, a list of 760 proteins that non-specifically bind to this tag or the associated purification beads has been assembled from data on 543 GS-based TAP experiments (Van Leene et al., 2015). We fused the GS^rhino^ TAP-tag to either the N-terminal end (*N-TAP-ARP6*) or the C-terminal end (*C-TAP-ARP6*) of the genomic ARP6 coding sequence and introduced the constructs into *arp6-1* mutant plants to test for the ability of each transgene to complement a null *arp6* allele. Using western blotting, we first detected the presence of the 67 kDa ARP6-TAP-tag fusion protein specifically in the plants homozygous for the transgene and not in the *arp6-1* or wild-type (WT) plants (Figure 1A). Next, we assessed the ability of the transgenes to rescue the morphological defects of *arp6-1* plants. When grown next to each other, the transgenic plants appeared almost indistinguishable from WT plants, with more compacted, non-serrated rosette leaves as compared to *arp6* mutants (Figure 1B). At the time of bolting, the average number of rosette leaves in wild type and transgenic plants was significantly higher than in *arp6-1* plants (Figure 1C), indicating that the *N-TAP-ARP6* and, to a lesser degree, the *C-TAP-ARP6* transgenes are able to rescue the early flowering phenotype of *arp6* plants (Figure 1D). Finally, all transgenic plants showed full complementation of the loss of apical dominance and fertility defects of *arp6-1* mutant plants (Figure 1E). Overall, we conclude that the *N-TAP-ARP6* and *C-TAP-ARP6* transgenes are fully functional, and thus suitable for affinity purification, since they were able to rescue the *arp6* mutant phenotypes.

**Figure 1.**
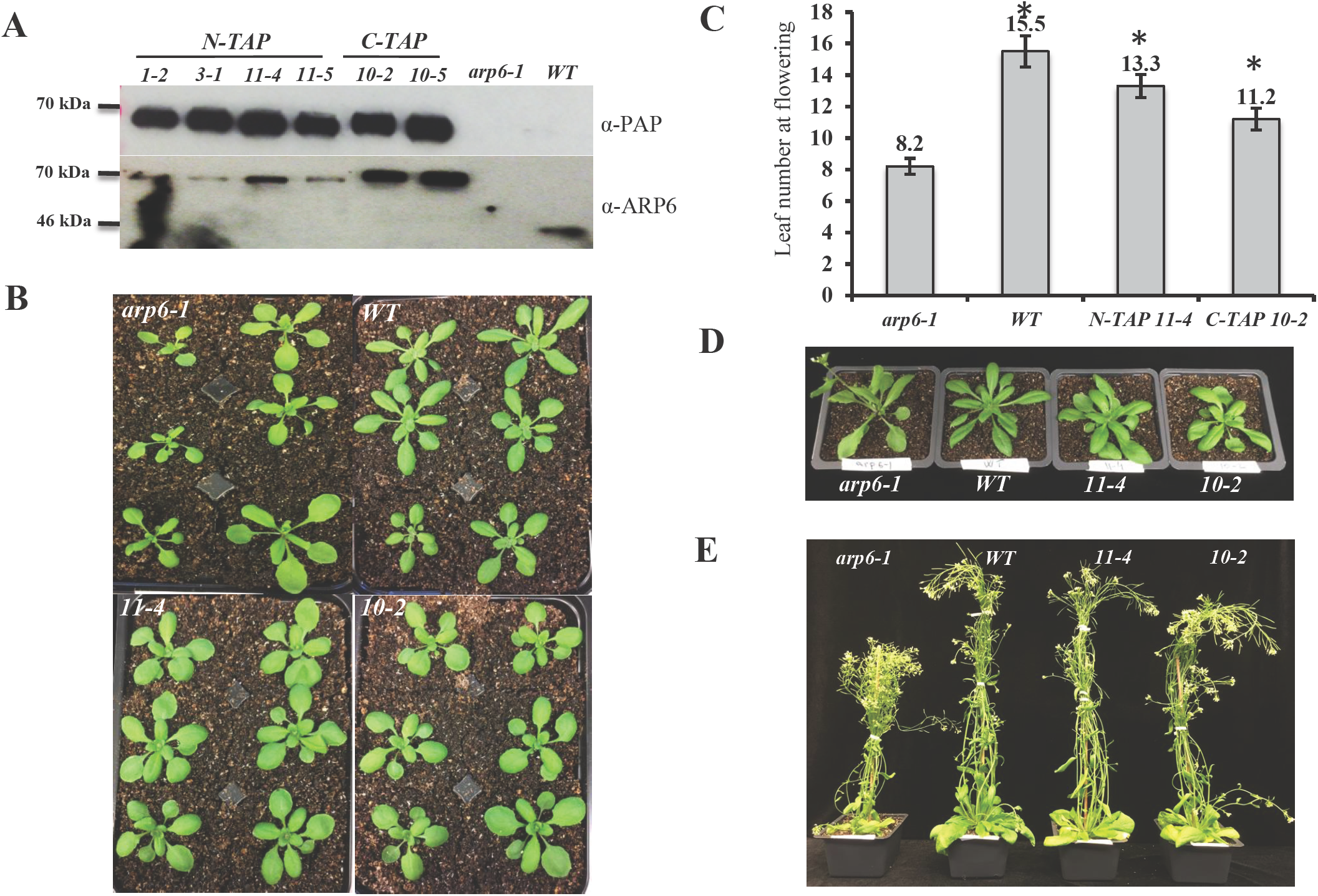
*N-TAP-ARP6* and *C-TAP-ARP6* transgenes rescue the *arp6-1* phenotype. (A-upper blot) T2 plants homozygous for *N-TAP-* or *C-TAP-ARP6* transgenes express a fusion protein with the expected size of 67.5 kDa. The fusion protein is specifically detected only in transgenic plants and not in *arp6-1* or WT plants using a peroxidase anti-peroxidase (PAP) soluble complex. (A-lower blot) The same protein extracts as in the upper blot were probed with a monoclonal ARP6 antibody. ARP6 presence is specifically detected in all transgenic plants as a higher, 67.5 kDa fusion protein band compared to the 44 kDa ARP6 band in WT plants, and is absent in *arp6-1* mutant plants. The ARP6 antibody reacts less strongly with N-TAP-ARP6, most likely because the antibody recognizes the N-terminal region which is adjoined to the TAP tag in this fusion. (B) Transgenic plants look more similar to the WT plants than *arp6-1* plants with more compacted, non-serrated rosette leaves. (C) The average number of rosette leaves of *N-TAP-ARP6* and *C-TAP-ARP6* transgenic plants at flowering is significantly higher than in *arp6-1* (n=6 for WT and *arp6-1*, and n=12 for *N-TAP 11-4* and *C-TAP 10-2*). Asterisks indicate significant differences from *arp6-1* plants with p<0.001, calculated using unpaired t tests. (D) Early flowering phenotype is rescued in transgenic plants when compared to the *arp6-1*. (E) The loss of apical dominance and fertility defects of *arp6-1* plants are completely rescued in *N-TAP-ARP6* and *C-TAP-ARP6* transgenic plants.

### Affinity purifications of ARP6-TAP-tag protein co-purified known components of the SWR1 complex and additional proteins

Since the *N-TAP-ARP6* and *C-TAP-ARP6* transgenes were fully functional, we proceeded with the affinity purification experiments using two independent *N-TAP-ARP6* transgenic lines (*N-TAP 11-4* and *N-TAP 1-2*) and one *C-TAP-ARP6* line (*C-TAP 10-2*). We followed the protocol described by Van Leene and colleagues (2015) to purify and elute the ARP6-TAP-tag interacting proteins and identified the eluted proteins by liquid chromatography coupled with tandem mass spectrometry (LC-MS/MS). All eluted proteins detected in our three TAP-tag experiments are listed in Supplemental Dataset 1. Using the database of non-specific binders of the GS^rhino^ TAP-tag, we eliminated many proteins from this list as false positives and compiled a list of reproducible ARP6- interacting proteins. Among these proteins, we identified ARP6, SWC2, SWC4, SWC6, PIE1, RuvB1, RuvB2, ACTIN1, ARP4, YAF9a, and H2A.Z proteins as Arabidopsis homologs of all 11 conserved subunits found in both yeast SWR1 and mammalian SRCAP complexes (Table 1). While we were able to detect either HTA9 or HTA11 in each of our three TAP-tag experiments, we never detected HTA8, the third member of the H2A.Z family. This finding is perhaps not surprising considering the fact that HTA8 is expressed at a very low level in Arabidopsis (Figure S1A). We also did not detect the YAF9b protein (Table 1) even though Arabidopsis YAF9a and YAF9b have been previously shown to act redundantly and are required for proper *FLC* expression (Bieluszewski et al., 2015; Choi et al., 2011). In addition to H2A.Z, we identified three H2B histones in our TAP experiments: HTB2, HTB4, and HTB9 (Table1). Since H2A.Z histones are deposited into nucleosomes as H2A.Z/H2B dimers, we sought to investigate whether specific H2A.Z proteins might have preferential H2B partners. If this is true, we would expect to see highly synchronized expression of specific H2A.Z/H2B pairs in various Arabidopsis tissues. Using publicly available microarray expression data (Schmid et al., 2005) for the two H2A.Z and three H2B histones that we identified, we observed that HTA11 and HTB2 had highly similar expression profiles across tissues (Figure S1B), while the expression of HTA9 matched very well with HTB4 expression and, to a slightly lesser degree, with HTB9 expression (Figure S1C and D). These results indicate that the Arabidopsis H2A.Z histones may have preferential H2B partners when deposited as dimers into nucleosomes.

**Table 1.** *Arabidopsis thaliana* proteins that co-purify with N-TAP ARP6 and C-TAP ARP6. We used mass spectrometry (MS) analysis to identify eluted proteins co-purified with ARP6-TAP-tag. All proteins identified by MS from three independent experiments (Table S1) were first filtered against the proteins purified from WT by TAP and then against the list of 760 known non-specific binders of the GS-TAP-tag. Table 1 contains the proteins that are not found on the list of the TAP-tag background proteins and are known homologs of SWR1 complex subunits in yeast and SRCAP subunits in mammals. The table also includes proteins such as MBD9, TRA1, and Alfins (lower rows in italics) that may represent novel subunits of the SWR1 complex. The table shows the expected MWs for each identified protein, and the total number of peptides identified by MS from two *N-TAP-ARP6* and one *C-TAP-ARP6* transgenic plants. ND=not detected.

| **Protein** | **Gene ID** | **Predicted MW (kDA)** | **Peptide number (N-TAP 11-4)** | **Peptide number (N-TAP 1-2)** | **Peptide number (C-TAP 10-2)** |
| --- | --- | --- | --- | --- | --- |
| **ARP6** | **AT3G33520** | **46.9** | **34** | **35** | **28** |
| **PIE1** | **AT3G12810** | **234** | **28** | **12** | **31** |
| **SWC2** | **AT2G36740** | **42.1** | **5** | **3** | **3** |
| **SWC4** | **AT2G47210** | **49.8** | **7** | **5** | **5** |
| **SWC6** | **AT5G37055** | **19.1** | **2** | **3** | **3** |
| **RuvB1** | **AT5G22330** | **50.3** | **25** | **22** | **21** |
| **RuvB2** | **AT5G67630** | **52.1** | **23** | **22** | **23** |
| **HTA8** | **AT2G38810** | **14.4** | **ND** | **ND** | **ND** |
| **HTA9** | **AT1G52740** | **14.3** | **2** | **ND** | **ND** |
| **HTA11** | **AT3G54560** | **14.5** | **ND** | **1** | **3** |
| **HTB2** | **AT5G22880** | **15.7** | **6** | **ND** | **5** |
| **HTB4** | **AT5G59910** | **16.4** | **8** | **5** | **5** |
| **HTB9** | **AT3G45980** | **16.4** | **8** | **ND** | **5** |
| **ARP4** | **AT1G18450** | **48.9** | **14** | **11** | **13** |
| **ACT1** | **AT2G37620** | **41.8** | **9** | **9** | **8** |
| **YAF9a** | **AT5G45600** | **30.2** | **1** | **ND** | **6** |
| **YAF9b** | **AT2G18000** | **30.4** | **ND** | **ND** | **ND** |
| **MBD9** | **AT3G01460** | **240.4** | **12** | **4** | **6** |
| **TRA1** | **AT4G36080** | **433.6** | **17** | **1** | **11** |
| **AL5** | **AT5G20510** | **29.6** | **3** | **2** | **2** |
| **AL6** | **AT2G02470** | **28.8** | **4** | **2** | **2** |
| **AL7** | **AT1G14510** | **28.3** | **3** | **ND** | **3** |

Interestingly, in addition to known subunits of the SWR1 complex, we also identified several nuclear proteins that were not previously reported as being associated with the SWR1 complex (Table 1). These include MBD9, a protein with a methyl-CpG-binding domain and various chromatin-binding domains (Aravind and Iyer, 2012; Peng et al., 2006; Yaish et al., 2009), TRA1, an uncharacterized Arabidopsis homolog of the NuA4 subunit Tra1 in yeast and the TIP60 subunit TRRAP in mammals (Bieluszewski et al., 2015; Lu et al., 2009), and three members of a plant-specific Alfin1-like family (AL5, AL6, and AL7) best known for their regulation of abiotic stress responses in Arabidopsis and ability to bind di- and tri-methylated lysine 4 of histone H3 (H3K4me2/3) (Wei et al., 2015).

In summary, we successfully purified the Arabidopsis SWR1 complex using the ARP6-TAP-tag as bait and conclude that the Arabidopsis SWR1 is a functionally and structurally conserved complex composed of the same 11 subunits found in the yeast SWR1 and human SRCAP complexes.

### MBD9 is required for H2A.Z incorporation into chromatin

One of the proteins identified in all three TAP-tag experiments as an ARP6-interacting partner was MBD9, a methyl-CpG-binding domain-containing protein. Previous studies have shown that *mbd9* mutants flowered significantly earlier than WT plants, due to reduced *FLC* expression, and produced more inflorescence branches when compared to WT plants (Peng et al., 2006), which are phenotypes also found in *arp6* mutants (Deal et al., 2005). We discovered that, in addition to the above-mentioned defects, *mbd9* plants have serrated rosette leaves and a significantly increased number of flowers with extra petals (Figure S2), which are phenotypes also associated with the loss of *ARP6* (Choi et al., 2005; Choi et al., 2007; Deal et al., 2005). Furthermore, examination of the previously reported MBD9 enrichment pattern at the *FLC* locus revealed that the two *FLC* regions with the highest H2A.Z enrichment in WT plants were also occupied by MBD9 (Deal et al., 2007; Yaish et al., 2009). Given that *arp6* and *mbd9* plants have similar phenotypes, that H2A.Z and MBD9 appear to occupy the same *FLC* regions, and that MBD9 co-purified with ARP6 in our TAP-tag experiments, we investigated whether MBD9 plays any role in the incorporation of H2A.Z into chromatin. We performed three biological replicates of chromatin immunoprecipitation coupled with high-throughput sequencing (ChIP-seq) using an H2A.Z antibody on WT, *arp6-1*, and *mbd9-1* seedlings. The average H2A.Z enrichment profile across all gene bodies in WT plants showed the highest enrichment of H2A.Z just after the transcription start site, with decreasing enrichment toward the 3’ end, as expected (Figure 2A). The pattern of H2A.Z enrichment across genes in *arp6-1* showed a similar profile but with extremely reduced enrichment, while *mbd9-1* plants had an intermediate level of H2A.Z enrichment between WT and *arp6* (Figure 2A). To analyze all of the regions normally enriched with H2A.Z, we identified peaks of H2A.Z enrichment that were present in at least two of the three of H2A.Z ChIP-seq replicates in WT plants and examined H2A.Z levels at these sites in *arp6-1* and *mbd9-1* mutants. As observed for gene bodies, the 7039 sites reproducibly enriched for H2A.Z in WT were nearly depleted of H2A.Z in *arp6-1*, while there was an intermediate H2A.Z enrichment level in *mbd9-1* plants (Figure 2B). These results indicate that MBD9 is indeed required for proper H2A.Z incorporation into chromatin. Since the loss of H2A.Z in *mbd9-1* plants is not as dramatic as in *arp6-1* plants on average (Figure 2A and B), MBD9 may be required for incorporation of H2A.Z at a subset of H2A.Z-enriched regions or may be required for full H2A.Z deposition at most sites.

**Figure 2.**
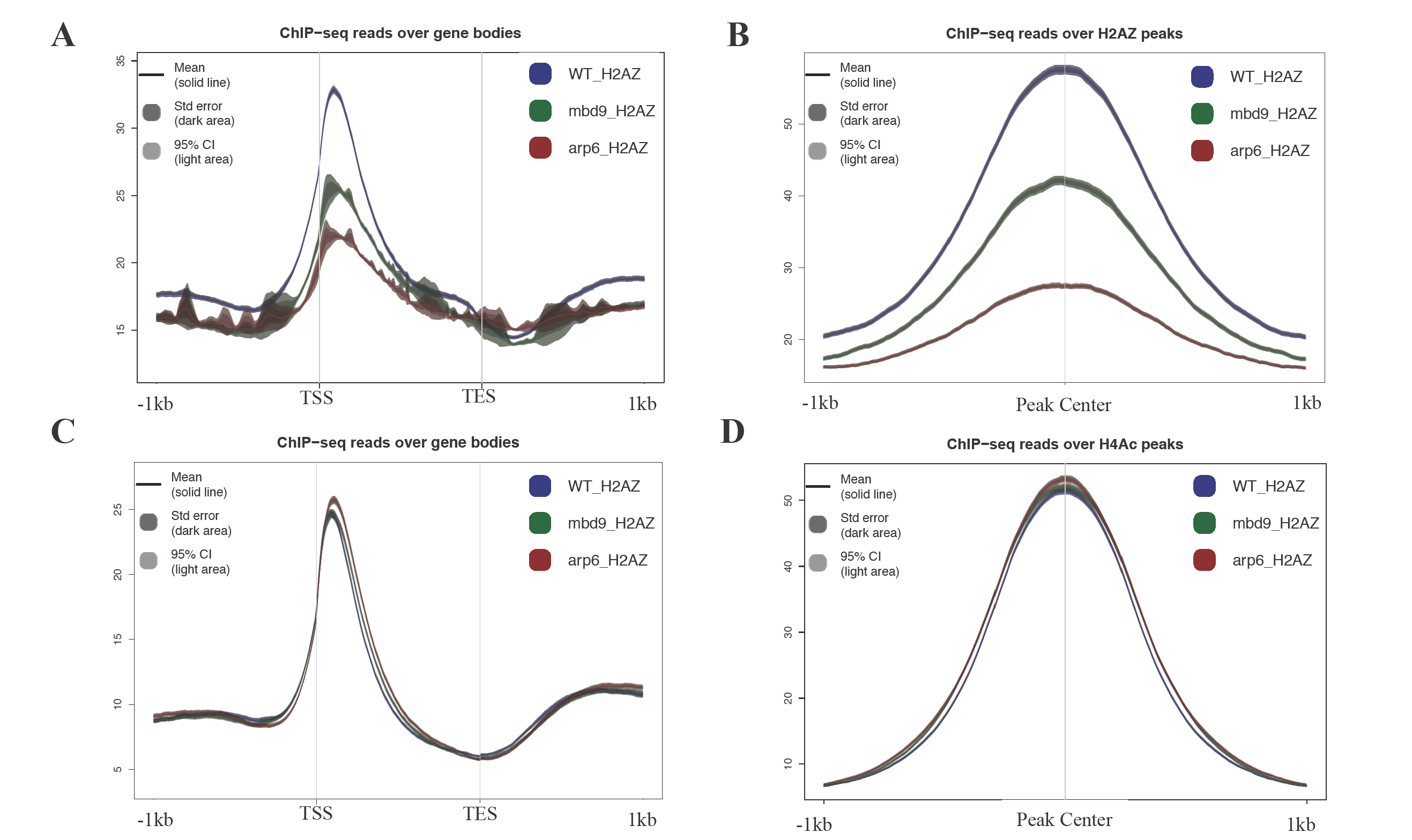
MBD9 is required for proper H2A.Z incorporation into chromatin but is not required for acetylation of H4. All ChIP-seq profiles of WT (shown in blue), *mbd9-1* (shown in green), and *arp6-1* (shown in red) were produced using SeqPlots. Standard error is represented as shaded divergence from the solid line of the mean signal. (A) Average ChIP-seq H2A.Z profiles plotted over gene body coordinates for all Arabidopsis genes, from the transcript start site (TSS) to the transcript end site (TES). (B) H2A.Z signal for each genotype over reproducible H2A.Z-enriched regions from WT plants. (C) Average ChIP-seq H4ac profiles plotted across gene bodies. (D) H4Ac signals over over reproducible H4Ac-enriched regions from WT plants.

As MBD9 was previously reported to have HAT activity and was found to associate with acetylated H4 (Yaish et al., 2009), we also examined the global level of histone H4 N-terminal acetylation in WT, *mbd9-1*, and *arp6-1* plants. If MBD9 is responsible for acetylation of H4 we would expect to see reduction of acetylated H4 in *mbd9-1* mutants compared to WT. However, we found that the genome-wide distribution of acetylated H4 was indistinguishable between WT, *mbd9-1*, and *arp6-1* plants when examined both across all gene bodies and at all sites enriched for acetylated H4 in WT plants (Figure 2C and D). This indicates that MBD9 does not globally affect H4 acetylation, and more importantly, these data also suggest that mutation of *MBD9* does not affect global nucleosome occupancy or modification per se, but specifically affects H2A.Z incorporation.

To confirm our results with respect to the role of MBD9 in H2A.Z deposition, we performed ChIP-qPCR experiments using WT, *arp6-1, mbd9-1*, and two additional *mbd9* T-DNA alleles *(mbd9-*2 *and mbd9-*3; Peng et al., 2006). We first assayed H2A.Z abundance at two distinct regions of the *FLC* gene: the first and last exon (regions 2 and 9, respectively, as described in Deal et al., 2007). Regions 2 and 9 are the sites on the *FLC* gene where H2A.Z is most highly enriched in WT plants, and that enrichment is lost in *arp6-1* mutant plants (Deal et al, 2007, Figure S3A). We found that in plants homozygous for any of the three *mbd9* alleles, the amount of H2A.Z at *FLC* regions 2 and 9 was reduced at least 2-fold when compared to WT plants (Figure S3A), indicating that MBD9 contributes to H2A.Z deposition at the *FLC* gene. To examine whether MBD9 regulates H2A.Z exchange at genes other than *FLC*, we measured the H2A.Z abundance at *ASK11* and *At4*, two phosphate starvation response genes previously shown to have H2A.Z deposited in their chromatin (Smith et al., 2010). We discovered that in *mbd9* plants these genes were depleted of H2A.Z to similar levels as in *arp6-1* plants when compared to the WT (Figure S3B). Taken together, our results indicate that MBD9 is required for H2A.Z deposition at multiple Arabidopsis genomic loci and is, therefore, functionally related to the SWR1 complex.

### A subset of H2A.Z-enriched sites require MBD9 for H2A.Z incorporation

To identify genomic sites that require MBD9 for H2A.Z incorporation into chromatin, we quantified the H2A.Z ChIP-seq reads from WT, *arp6-1*, and *mbd9-1* mutant plants across all of the H2A.Z-enriched regions that were reproducibly identified in the WT replicates. We then performed DESeq analysis (Anders and Huber, 2010) to quantitatively compare WT to *mbd9-1* and WT to *arp6-1* H2A.Z levels at each site (Supplemental Dataset 2). Out of the total of 7039 H2A.Z-enriched sites, we identified 1391 H2A.Z sites that were reproducibly reduced at least 1.5-fold (log fold change of at least 0.6 with an adjusted p value ≤ 0.05) in *mbd9-1* compared to WT (Figure 3A). In contrast, H2A.Z levels were significantly depleted in *arp6-1* at nearly all of the H2A.Z sites, as expected for a mutation that disrupts the SWR1 complex (Figure 3B).

**Figure 3.**
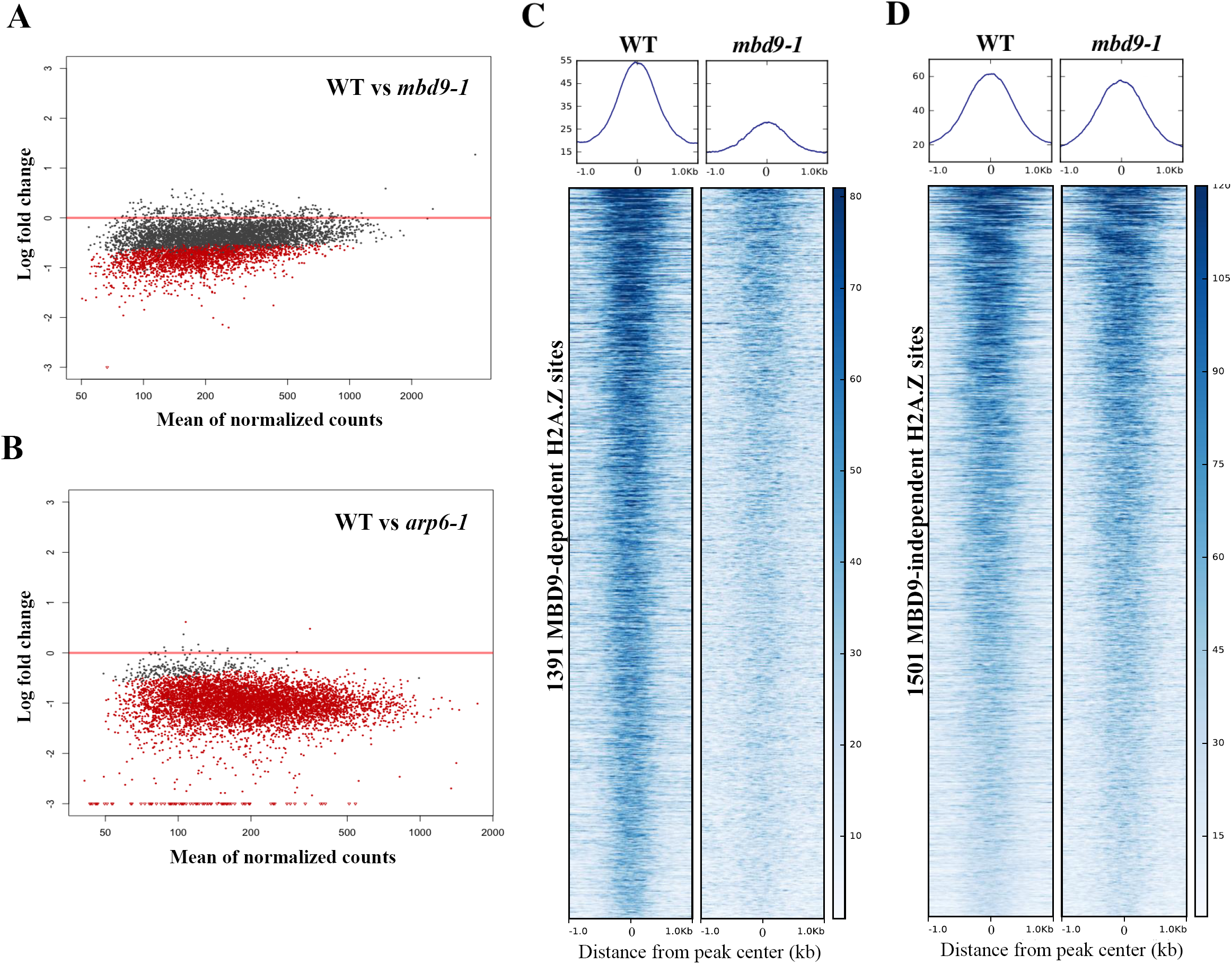
Identification of 1391 H2A.Z-enriched sites that require MBD9 for H2A.Z incorporation into chromatin. (A) and (B) Scatter plots of log fold changes of H2A.Z ChIP-seq reads (y-axis) versus mean of normalized read counts (X-axis) between wild type plants and the indicated mutants. Each dot represents the log fold change for one reproducible H2A.Z-enriched peak region identified in WT plants. Dots in red indicate a statistically significant log fold change between genotypes for that peak. (A) H2A.Z read count comparison between WT and *mbd9-1* H2A.Z levels. Using the cutoff of log fold change of at least 0.6 with an adjusted p value of 0.05, there are 1391 sites that are significantly depleted for H2A.Z in *mbd9-1* plants. (B) H2A.Z read counts compared at the same peak coordinates as in (A), but between WT and *arp6-1* plants. (C) and (D) Average plots and heatmaps of the 1391 MBD9-dependent (C) and 1505 MBD9-independent H2A.Z peaks (D), centered on the peak.

To further examine the nature of the H2A.Z deposition defect in *mbd9* mutants, we visualized H2A.Z enrichment and distribution across the 1391 sites that lose H2A.Z in *mbd9-1*, which we refer to as MBD9-dependent H2A.Z sites. For comparison, we selected an equivalently-sized set of MBD9-independent H2A.Z sites (1505 sites with an average fold difference of less than 1.19 between WT and *mbd9-1*, which is an absolute log fold change of less than 0.25). This analysis revealed a drastic reduction in H2A.Z occupancy at each of the MBD9-dependent H2A.Z sites when comparing WT and *mbd9-1*, but with maintenance of the same overall pattern of occupancy (Figure 3C). In contrast, the MBD9-independent H2A.Z sites showed equivalent profiles and occupancy levels between WT and *mbd9-1* (Figure 3D). Thus, MBD9 is required for proper H2A.Z deposition at a subset of H2A.Z sites and may act through recruitment of the SWR1 complex to these sites.

### MBD9-dependent H2A.Z sites have chromatin features distinct from the MBD9- independent H2A.Z sites

In order to understand why MBD9 is required for H2A.Z deposition at certain sites and not others, we first analyzed the genomic distribution of MBD9-dependent H2A.Z sites compared to our size-matched set of MBD9-independent H2A.Z sites. We found that the two sets are distributed similarly across the genome, with more than 80% of each set of coordinates localizing within genic regions (Supplemental Figure S4). Next, we identified the 1322 genes associated with MBD9-dependent H2A.Z sites (Supplemental Dataset 2) and performed Gene Ontology (GO) enrichment analysis. However, no significantly overrepresented GO terms were identified using either of two different GO analysis tools, indicating that MBD9-mediated deposition of H2A.Z is not detectably associated with functionally-related gene sets or particular cellular pathways.

We also examined various histone modification profiles at the two types of sites using publicly available ChIP-seq data from WT plants, in order to discern any differences between H2A.Z sites that require MBD9 and those that do not. Interestingly, we found that in WT plants the average level of acetylation of histone H3 at lysine 9 (H3K9Ac) is higher at MBD9-dependent H2A.Z sites than it is at the sites that do not require MBD9 (Figure 4A). However, no differences were found in the average enrichment of acetylation of histone H3 at lysine 18 and 27 (H3K18Ac and H3K27Ac, respectively) between the two types of loci (Figure 4B and C). On the other hand, the enrichment of di-methylation at lysine 9 of histone H3 (H3K9me2) reads was reversed compared to the H3K9Ac enrichment levels at these two types of loci, being less abundant at MBD9-dependent H2A.Z sites (Figure 4D). Furthermore, the average ChIP signals in WT plants for histone H3 trimethylation at lysine 4 or lysine 36 (H3K4me3 and H3K36me3, respectively) were also consistently lower at the MBD9-dependent H2A.Z sites (Figure 4E and F). Collectively, the sites that require MBD9 for H2A.Z incorporation appear to have several distinct chromatin features, including higher levels of H3K9Ac as well as a depletion of H3K4me3 and H3K36me3, compared to the sites that do not require MBD9.

**Figure 4.**
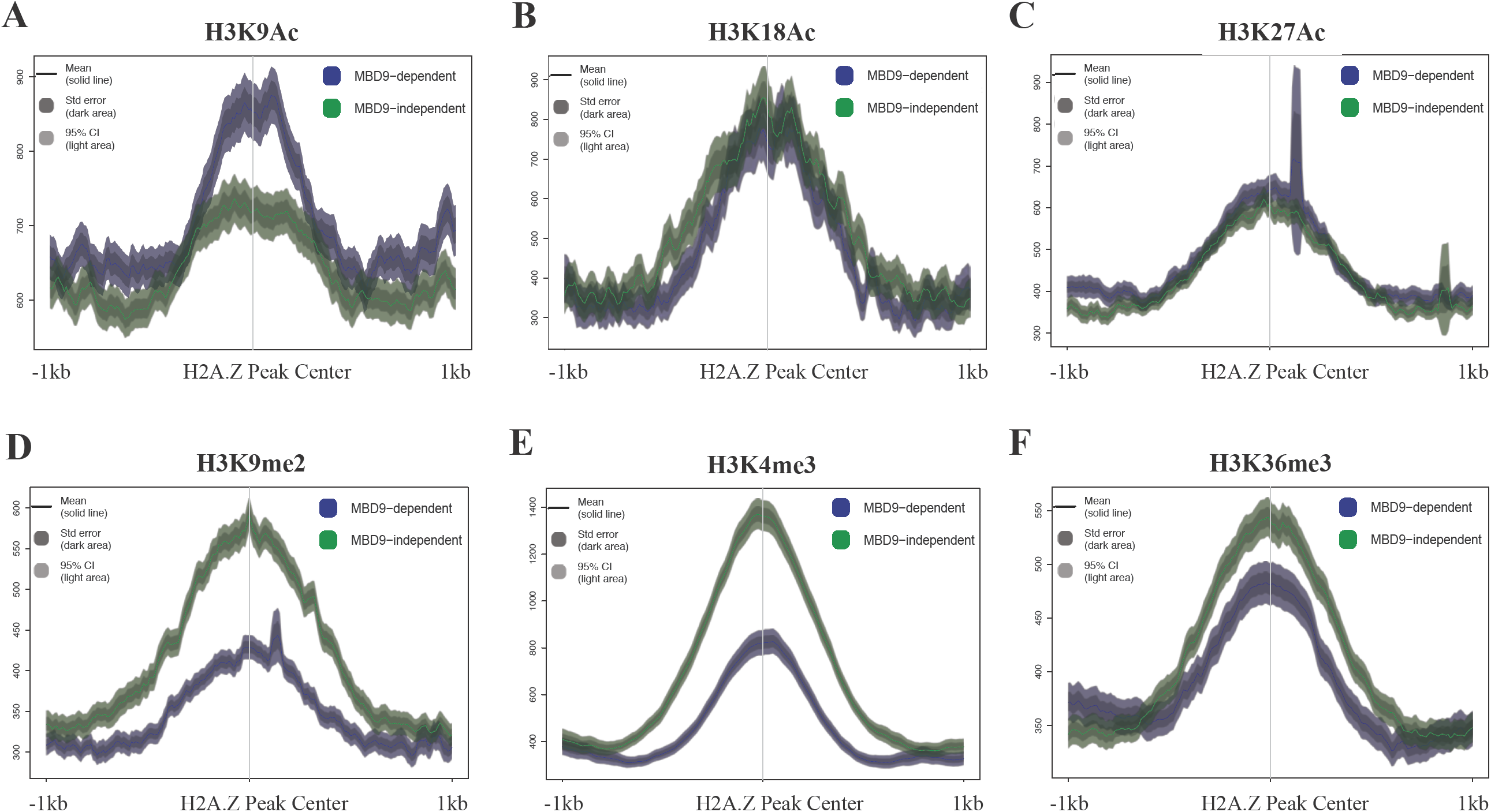
H2A.Z sites that require MBD9 have distinct chromatin properties. Average ChIP-seq profiles of H3K9Ac (A), H3K18Ac (B), H3K27Ac (C), H3K9me2 (D), H3K4me3 (E), and H3K36me3 (F) at MBD9-dependent H2A.Z sites (shown in blue) and MBD9-independent H2A.Z sites (shown in green). MBD9-dependent H2A.Z sites appear to have several distinct chromatin features, including higher levels of H3K9Ac (A), as well as a depletion of H3K4me2 (D), H3K4me3 (E), and H3K36me3 (F), compared to the sites that do not require MBD9.

Given the predominant gene-body localization of H2A.Z, the differences in histone modification levels between MBD9-dependent and-independent sites could simply reflect differences in expression levels of the underlying genes. However, we found that genes nearest to the sites in each category span a wide range of expression levels and are not significantly different from one another in terms of steady-state transcript levels (unpaired t-test, p < 0.05, Supplemental Figure 5). Thus, MBD9 may recognize specific chromatin features, for example the H3K9Ac mark via its bromodomain (Yang, 2004), which could help guide the SWR1 complex to specific DNA sites.

**Figure 5.**
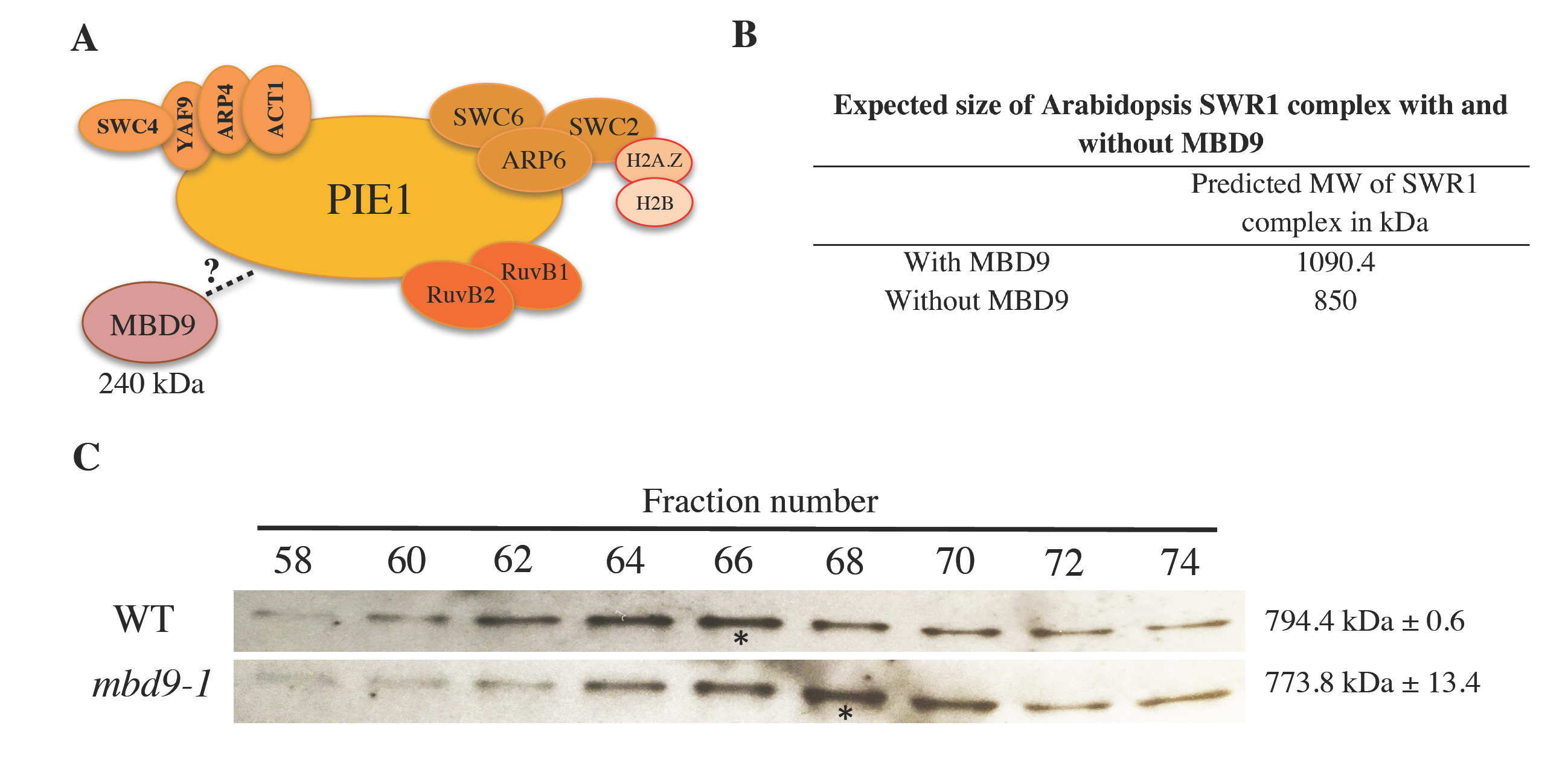
MBD9 is not a stable subunit of the Arabidopsis SWR1 complex. (A) Schematic representation of the Arabidopsis SWR1 protein complex based on TAP-MS experiments, including MBD9 as a potential SWR1 subunit. (B) Estimated size of the Arabidopsis SWR1 complex with and without MBD9 as a subunit. The estimated size was calculated based on the known stoichiometry of the yeast SWR1 complex (Nguyen et al., 2013) and predicted molecular weights of Arabidopsis SWR1 subunits listed in Table 1. (C) Protein gel blots of even-numbered SEC fractions from WT (top blot) and *mbd9-1* (bottom blot) plants. The blots were incubated with the ARP6 monoclonal antibody (Deal et al., 2005). Asterisks indicate the ARP6 peak fractions. The average molecular weights of ARP6-containing protein complexes in WT and *mbd9-1* plants were calculated from two biological replicates and presented on the right side of the corresponding protein blots.

### MBD9 is not a core subunit of the Arabidopsis SWR1 complex

To determine whether the MBD9 protein, with an estimated molecular mass of 240 kDa, is an integral component of the Arabidopsis SWR1 complex (Figure 5A) we performed size-exclusion chromatography (SEC) experiments on protein extracts from WT and *mbd9-1* plants, followed by western blotting for ARP6. This allowed us to define the native size of the complex and to determine whether this size changes in the absence of MBD9, as would be expected if this protein were a stoichiometric component of the SWR1 complex (Figure 5B), as previously demonstrated for the PIE1 subunit (Deal et al., 2007). Using an ARP6 monoclonal antibody (Deal et al., 2005), we detected ARP6 protein in its native form as a part of a multi-subunit complex with a molecular mass of ∼800 kDa (Figure 5C). When the SEC experiments were performed on *mbd9-1* extracts, the ARP6 peak did not significantly shift and the estimated molecular mass of the native ARP6 complex in *mbd9-1* plants was ∼775 kDa (Figure 5C) in two biological replicates. These results strongly suggest that MBD9 is not a core component of the ARP6- containing SWR1 complex, but most likely interacts with it in a more transient manner.

### arp6-1;mbd9-1 double mutant plants have a more severe phenotype than either single mutant

So far, we discovered that MBD9 is functionally related to, but not stably associated with, the Arabidopsis SWR1 complex. To investigate genetic interactions between *MBD9* and *ARP6* we generated *arp6-1;mbd9-1* double mutant plants. We have shown that single *arp6-1* and *mbd9-1* mutant plants have similar phenotypic defects (Figure S2) and that both ARP6 and MBD9 regulate H2A.Z incorporation into chromatin (Figure 2A and B). If these two proteins are subunits of the same complex or function exclusively in the same genetic pathway then double mutant plants should be phenotypically indistinguishable from single mutants, as previously shown for *arp6;swc6* plants (Choi et al., 2007; Lazaro et al., 2008). Instead, we observed that the double mutants displayed much more severe defects (dwarf stature, deformed leaves, and drastically reduced fertility) than the individual single mutants throughout development (Figure 6). Importantly, these phenotypes in the double mutant plants were reverted back to those of each single mutant by introducing either the genomic *ARP6* or *MBD9* constructs into the double mutants (Figure S6), indicating that these defects were truly the result of simultaneous loss of ARP6 and MBD9 functions. These findings further support the idea that MBD9 is not a core subunit of the ARP6-containing SWR1 complex and suggest that this protein has additional functions outside of H2A.Z incorporation (Hale et al., 2016).

**Figure 6.**
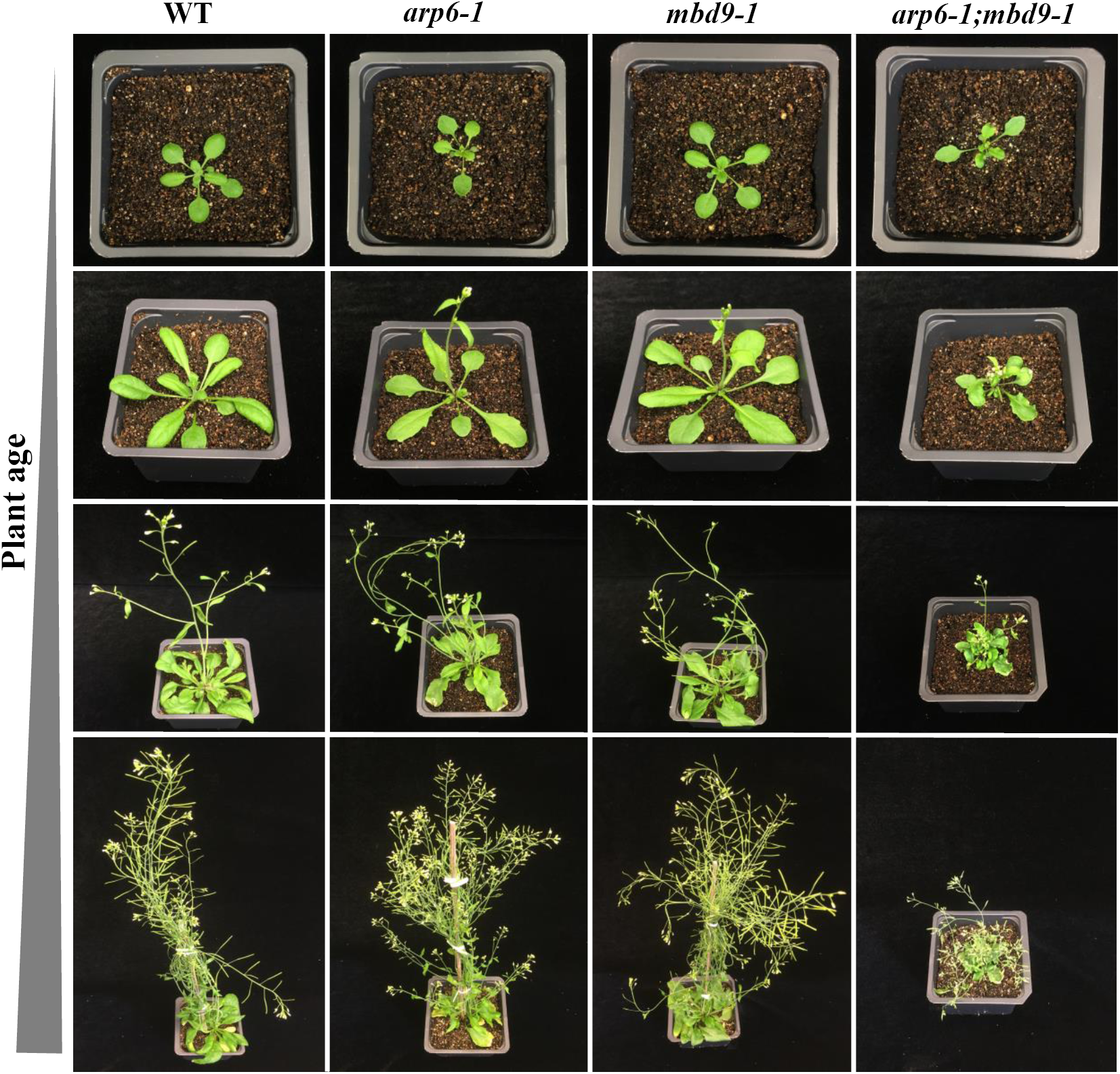
*arp6-1;mbd9-1* double mutant plants have a more severe phenotype than either single mutant. WT, *arp6-1, mbd9-1*, and *arp6-1;mbd9-1* plants grown under long-day conditions were individually photographed at four time points over five weeks of growth. *arp6-1;mbd9-1* double mutant plants have severely delayed development and are dwarfed compared to single *arp6-1* and *mbd9-1* mutant plants.

## DISCUSSION

### The SWR1 complex is conserved in plants

Previous studies provided important but rather circumstantial evidence that Arabidopsis contains a SWR1 complex that mediates incorporation of H2AZ into chromatin (Bieluszewski et al., 2015; Choi et al., 2005; Choi et al., 2007; Deal et al., 2005; Deal et al., 2007; Gomez-Zambrano et al., 2018; Lazaro et al., 2008; March-Diaz et al., 2007; March-Diaz et al., 2008; March-Diaz and Reyes, 2009; Martin-Trillo et al., 2006; Noh and Amasino, 2003). Importantly, no previous study has described the subunit composition of this complex in plants. Using the SWR1-specific subunit ARP6 as bait, we successfully purified the Arabidopsis SWR1 complex and identified all 11 conserved subunits that are also found in the yeast SWR1 and mammalian SRCAP complexes. These results suggest that the function and structure of the canonical SWR1 complexes that incorporate histone H2A.Z into the nucleosomes have been well preserved over evolutionary timescales and may be found in all eukaryotes.

It has been previously shown that SWC4, ARP4, YAF9, and ACT1 are subunits shared between yeast SWR1 and NuA4 complexes (Kobor et al., 2004; Krogan et al., 2003; Lu et al., 2009; Mizuguchi et al., 2004; Zhang et al., 2004). The results of Bieluszewski and colleagues suggest that these four subunits may also be shared between Arabidopsis SWR1 and NuA4 complexes (Bieluszewski et al., 2015). They used SWC4 and ARP4 as baits to purify many proteins homologous to the yeast SWR1 and NuA4 complex components, including Arabidopsis EAF1, which in yeast serves as a platform subunit of the NuA4 complex (Auger et al., 2008). The identification of an Arabidopsis EAF1 subunit strongly implies that plants possess an independent NuA4 complex. However, to unequivocally confirm this, EAF1 would need to be used as bait to purify the EAF1-containing protein complex alone.

SWC4, ARP4, Yaf9, and ACT1 are also shared subunits between mammalian SRCAP and TIP60 complexes (Lu et al., 2009). As mentioned earlier, TIP60 in higher eukaryotes is a single multifunctional complex that combines the subunits and functions of yeast SWR1 and NuA4 complexes (Lu et al., 2009). It appears that this merger evolved as a result of the fusion of the two major scaffolding proteins, the Swr1 ATPase of the yeast SWR1 complex and Eaf1 protein of the yeast NuA4 complex, into a single p400/Domino-like protein. This is based on the fact that p400/Domino contains HSA, ATPase, and SANT domains, which are found separately in the Swr1 and Eaf1 proteins (Bieluszewski et al., 2015; Lu et al., 2009). As a result of this protein fusion event, the genomes ancestral to Drosophila and mammals lost Eaf1 homologs and therefore do not possess canonical NuA4 complexes (Lu et al., 2009). In Arabidopsis, PIE1 is a component of the SWR1 complex and acts as a homolog of the yeast Swr1. Intriguingly, PIE1 also contains HSA, ATPase, and SANT domains, implying that PIE1 may also be an ortholog of p400/Domino. In fact, Bieuszewski and colleagues originally hypothesized that PIE1 is a scaffolding component of an Arabidopsis TIP60-like complex (Bieluszewski et al., 2015). Even though they subsequently purified the Arabidopsis EAF1 subunit and suggested that plants most likely possess an independent NuA4 complex, the authors did not exclude the possibility that Arabidopsis may still contain a TIP60-like complex organized around PIE1 (Bieluszewski et al., 2015). Interestingly, one of the proteins that co-purified with ARP6 in our TAP experiments was the TRA1 protein, an uncharacterized Arabidopsis homolog of the yeast NuA4 subunit Tra1 and the mammalian TIP60 subunit TRRAP, further suggesting an intimate functional relationship that may exist among Arabidopsis SWR1 and NuA4 complexes (Table 1). Taken together, it is plausible that plants possess both the canonical SWR1 complex and an independent NuA4-like complex, as in yeast, and also contain a TIP60-like complex, which is found only in higher eukaryotes. Future purification experiments using Arabidopsis PIE1 as bait are crucial to address the question of whether plants have two distinct PIE1-containing complexes (SWR1 and a TIP60-like), and which subunits are shared between the two complexes. The existence of two distinct PIE1 complexes could also explain why the phenotype of *pie1* mutant plants is more severe than those of *h2a.z* mutants or mutations in other SWR1 components (Choi et al., 2005; Choi et al., 2007; Coleman-Derr and Zilberman, 2012; Deal et al., 2005; Deal et al., 2007; Lazaro et al., 2008; March-Diaz et al., 2007; Martin-Trillo et al., 2006; Noh and Amasino, 2003).

### MBD9 may recruit the SWR1 complex to mediate H2A.Z deposition to chromatin

Based on multiple studies in many model organisms, we now have a good understanding of how the SWR1 complex incorporates H2A.Z into nucleosomes (Luk et al., 2010; Mizuguchi et al., 2004; Nguyen et al., 2013). However, several aspects of SWR1 biology are still poorly understood, including precisely how the SWR1 complex is recruited to specific chromatin regions to deposit H2A.Z. In yeast, it has been shown that NuA4- mediated acetylation of specific histones in the nucleosome is important for SWR1 targeting to chromatin and H2A.Z incorporation (Altaf et al., 2010; Babiarz et al., 2006; Cheng et al., 2015; Keogh et al., 2006; Lu et al., 2009; Millar et al., 2006). In addition, it has been proposed that Bdf1, a bromodomain-containing subunit of the yeast SWR1 complex, recruits the complex to chromatin by recognizing acetylated H4 tails (Altaf et al., 2010; Ladurner et al., 2003). Supporting this notion, the loss of Bdf1 results in global reduction of H2A.Z in chromatin (Durant and Pugh, 2007).

In plants, little is known about the mechanisms that target the SWR1 complex to specific chromatin loci. Recent results from the Jarillo group suggest that the binding of the SWC4 subunit to AT-rich DNA elements in promoters of certain genes can recruit the Arabidopsis SWR1 complex to these chromatin regions to deposit H2A.Z (Gomez-Zambrano et al., 2018). However, only a subset of H2A.Z-enriched genes contain AT-rich elements in their promoters, which strongly suggests that additional mechanisms of SWR1 recruitment to chromatin exist in plants. What role may MBD9 play in this process? In addition to the methyl-CpG-binding (MBD) domain, MBD9 protein contains a single bromodomain, which is known to recognize acetylated histones (Yang, 2004), and two plant homeodomains (PHD), which may recognize methylated lysines in histone H3 (Li et al., 2006). The presence of these domains in MBD9 suggests that the protein likely interacts with a specific pattern of modified histones (Berg et al., 2003; Peng et al., 2006), and a previous study has indeed shown that MBD9 associates with acetylated histones (Yaish et al., 2009). We have demonstrated that in *mbd9* mutant plants the level of H2A.Z incorporation is significantly reduced at a subset of H2A.Z-enriched regions (Figure 3) and that these MBD9-dependent H2A.Z loci have distinct histone modification profiles relative to H2A.Z-enriched regions that do not depend on MBD9 (Figure 4). Specifically, H2A.Z sites that are dependent on MBD9 had higher levels of H3K9Ac and lower levels of H3K4me3 and H3K36me3. This suggests that MBD9, like Bdf1 in yeast, could target SWR1 via its bromodomain by recognizing acetylated histone marks, such as H3K9Ac, and may also recognize the methylation state of H3K4 and H3K36 via its two PHD domains.

Animal and plant proteins that contain MBD domains are known to bind to methylated CpGs (m^5^CpG) both *in vitro* and *in vivo* (Du et al., 2015; Scebba et al., 2003; Zemach and Grafi, 2007). It has been proposed, however, that Arabidopsis MBD9 is not able to recognize m^5^CpG since its MBD domain is missing several amino acid residues required for binding to methylated CpGs (Berg et al., 2003). In addition to the MBD domain, MBD9 also contains Bromo, PHD, DDT, and WHIM domains, giving it a domain architecture highly similar to that of the human BAZ2A and BAZ2B proteins (Aravind and Iyer, 2012; Du et al., 2015). The BAZ2A protein is a component of the nucleolar remodeling complex (NoRC), and it has been shown that its MBD domain can bind to unmethylated DNA (Strohner et al., 2001).

Previous studies have demonstrated that H2A.Z and DNA methylation are antagonistic chromatin marks, and that the presence of one is anti-correlated with that of the other (Conerly et al., 2010; Zemach et al., 2010; Zilberman et al., 2008). While DNA methylation globally inhibits H2A.Z incorporation into chromatin, H2A.Z presence can exclude DNA methylation (Zilberman et al., 2008). In fact, it has been observed that *pie1* mutant plants show a small but consistent increase in DNA methylation genome-wide (Coleman-Derr and Zilberman, 2012; Zilberman et al., 2008), a phenotype also found in *mbd9-1* plants (Yaish et al., 2009). Perhaps the inability of MBD9 to bind methylated CpGs (Berg et al., 2003), combined with the fact that MBD9 is structurally similar to BAZ2A protein, which is capable of binding to unmethylated DNA (Strohner et al., 2001), could at least partially account for the targeting of H2A.Z to the regions that are free of DNA methylation (Coleman-Derr and Zilberman, 2012; Zilberman et al., 2008). Together, the combination of Bromo, PHD, and MBD domains may target MBD9 (and thereby SWR1) to unmethylated genomic regions that have a specific combination of histone modifications, potentially explaining why MBD9 is only responsible for H2A.Z incorporation at certain sites and not others.

Although MBD9 was co-purified in all of our ARP6 TAP-tag experiments (Table 1), MBD9 appears not to be a core component of the SWR1 complex (Figure 5). Two possible conclusions about MBD9’s interaction with the SWR1 complex can be made based on these results. MBD9 may interact only transiently with components of the SWR1 complex, and is therefore detected in TAP-tag experiments as previously demonstrated for transcription factors and cofactors that recruit Arabidopsis SWI/SNF and PRC2 complexes to specific chromatin sites (Efroni et al., 2013; Vercruyssen et al., 2014; Wu et al., 2012; Xiao et al., 2017). Alternatively, MBD9 could be more tightly associated with only a subset of all SWR1 complexes in Arabidopsis. In that case, our size exclusion chromatography experiment most likely would not show a significant shift in *mbd9-1* plants when compared to WT because the loss of MBD9 affects only a minor fraction of SWR1 complexes. While further experiments are needed to determine the precise nature of MBD9’s interaction with the SWR1 complex, we show conclusively that MBD9 is functionally associated with the SWR1 complex and is integral to the deposition of H2A.Z at a subset of loci.

The presence of H2A.Z in chromatin has been linked to both gene activation and gene repression, but how H2A.Z affects transcription in this context-dependent manner is not clear (Surface et al., 2016; Zlatanova and Thakar, 2008). In addition, how the chromatin remodelers that deposit H2A.Z are recruited to specific chromatin loci is poorly understood. Our isolation of the Arabidopsis SWR1 complex identified unexpected proteins that co-purified with this complex, including MBD9 and three members of the plant-specific Alfin family. Based on our results and data from other studies, we propose that these SWR1-associated proteins may be involved in the recruitment of the SWR1 complex to chromatin to incorporate H2A.Z at specific loci in the Arabidopsis genome. With the identification of these proteins, we can now start to address important mechanistic questions about the activity of SWR1 in plants and how MBD9, and perhaps Alfins, may differentially modulate SWR1 functions during development.

## METHODS

### Plant material, growth conditions, and transformation

*Arabidopsis thaliana* of the Columbia (Col-0) ecotype was used as the wild type reference, and all mutant seeds are of the Col-0 ecotype. The *arp6-1* (SAIL_599_G03), and *mbd9-1* (SALK_054659), *mbd9-2* (SALK_121881) and *mbd9-3* (SALK_039302) alleles were described previously (Deal et al., 2005; Peng et al., 2006). Seedlings were grown in either soil, half-strength Murashige and Skoog (MS) liquid media (Murashige and Skoog, 1962), or on half-strength MS media agar plates, in growth chambers at 20°C under a 16 hour light/8 hour dark cycle. Plasmids containing *N-TAP-ARP6, C-TAP-ARP6*, and *gMBD9* constructs were introduced into the *Agrobacterium tumefaciens GV3101* strain by electroporation. Plants were transformed with these constructs via the floral dip method (Clough and Bent, 1998). Primary transgenic plants were selected on half-strength MS media agar plates containing 50 mg/L hygromycin and 100 mg/L timentin, and then transferred to soil. Two to three grams of sterilized WT seeds and *T*_*3*_ seeds homozygous *for N-TAP-ARP6* and *C-TAP-ARP6* constructs were germinated for 6 days in flasks containing 600 ml of half-strength MS media with constant shaking on rotating platform (80-90 rpm). After 6 days, the germinated seedlings were filtered to remove the excess liquid, and 50 grams of seedling tissue was frozen in liquid nitrogen and stored at −80°C.

### Plasmid DNA constructs

To construct ARP6-TAP-tag we fused genomic *ARP6* sequence to the tandem affinity purification (TAP) GS^rhino^ tag, recently developed for efficient affinity purifications of protein complexes in plants (Van Leene et al., 2015). Gateway–compatible plasmids containing either a C-terminal TAP-tag (pEN-R2-GS_rhino-L3, Van Leene et al., 2015) or an N-terminal TAP-tag (pEN-L1-NGS_rhino-L2, Van Leene et al., 2015) were used to produce the *C-TAP-ARP6* and the *N-TAP-ARP6* constructs, respectively. To generate the *C-TAP-ARP6* construct, a total of six primers were used in three overlapping PCR reactions to produce a ∼4.7 kb *attB* PCR fragment. This PCR product contained ∼4.1 kb of the genomic *ARP6* sequence (from −2040 bp upstream of the start codon to +2083 bp downstream from the start codon), ∼600 bp of the TAP-tag sequence fused to the C-terminal end of the *ARP6* gene, and *attB* adapters at 5’ and 3’ ends of the PCR product for Gateway cloning. This PCR fragment was sub-cloned into pDONR221 gateway plasmid via BP recombination reaction using BP clonase II enzyme (Invitrogen). The construct was verified by sequencing and further sub-cloned into the destination gateway plasmid pMDC99 (Curtis and Grossniklaus, 2003) using the LR clonase II enzyme in LR recombination reaction (Invitrogen). Similarly, the *attB N-TAP-ARP6* construct was first produced using six PCR primers in overlapping PCR reactions containing the same genomic *ARP6* DNA fragment as in the *C-TAP-ARP6*, with the TAP-tag fused at the N-terminal end of the *ARP6* gene. This PCR fragment was then sub-cloned into pDONR221 via BP reaction, verified by sequencing, and finally sub-cloned into the pMDC99 destination plasmid via LR reaction.

To generate *gMBD9* construct, which was used to transform *arp6-1;mbd9-1* double mutant plants, we first PCR-amplified 11,311 bp of genomic *MBD9* sequence (from –1936 bp upstream of the start codon to + 9372 downstream from the start codon) using *gMBD9* sequence-specific primers with *attB* adapters at their 5’ ends. The *attB* PCR product was then sub-cloned into pDONR221 gateway plasmid via BP recombination reaction (Invitrogen), verified by sequencing, and finally sub-cloned into the destination gateway plasmid pMDC99 (Curtis and Grossniklaus, 2003) using LR recombination reaction (Invitrogen).

### Crosses and genotyping

To produce *arp6-1;mbd9-1* double mutant plants, pollen from *arp6-1* plants was used to manually pollinate *mbd9-1* plants. Since *ARP6* and *MBD9* genes are both on chromosome 3, we were only able to identify the F_2_ plants that were homozygous for one T-DNA allele and heterozygous for the other. We used F_3_ seeds from *arp6-1/arp6-1;mbd9-1/+* plants to identify the double mutant plants.

### Protein gel blotting

The proteins for western blot detection were extracted from ∼100 mg of whole transgenic seedlings homozygous for the *arp6-1* allele and either the *C-TAP-ARP6* or *N-TAP-ARP6* constructs by first making a crude nuclei preparation using Nuclei Purification Buffer (Deal and Henikoff, 2010). The nuclei pellets were then resuspended in 2 volumes of 2x Laemmli’s sample buffer (125 mM Tris-HCl pH 6.8, 4% SDS, 30% glycerol, and 1% β- mercaptoethanol) prior to heating and loading on a gel. For ARP6 detection on fractions from SEC experiments (see below), the eluted proteins were isolated by adding 20 µl of the StrataClean resin (Agilent) to 1 ml of each SEC fraction, incubating for 20 minutes at room temperature (RT) on a rotating platform, and then spinning down for 2 minutes at 5,000g at RT. The pelleted proteins were resuspended in 20 µl of 2x Laemmli’s sample buffer. The proteins were then separated on 4-20% Novex WedgeWell tris-glycine gel (Invitrogen) and transferred to Amersham nitrocellulose blotting membrane (GE Healthcare). After blocking overnight in PBST buffer (137 mM NaCl, 2.7 mM KCl, 10 mM Na_2_HPO4, 2 mM KH_2_PO4, and 0.05% tween^®^20) containing 5% non-fat dry milk, the blots were incubated with primary antibody (1:100 dilution for monoclonal mouse ARP6 antibody (Deal et al., 2005), and 1:2,000 dilution for peroxidase anti-peroxidase (PAP) soluble complex antibody (Sigma-Aldrich) that detects the TAP-tag) in blocking solution for 1 hour at RT. The blots were washed 3 times for 5 minutes in PBST. The ARP6 blot was then incubated with the anti-mouse horseradish peroxidase-conjugated secondary antibody (1:2,000 dilution, GE Healthcare). The ARP6 blot was washed 3 more times for 5 minutes in PBST, and both blots were then incubated with ECL detection reagents for 3 minutes (Thermo Scientific) and used to expose Amersham Hyperfilm ECL (GE Healthcare) to detect protein bands.

### Purification of protein complexes containing the ARP6-TAP-tag fusion protein

ARP6-TAP-containing protein complex was purified as described in Van Leene et al., 2015, with following modifications: 1) instead of using a kitchen blender in a stainless steel wine cooler, 50 grams of frozen seedlings were ground with mortar and pestle in liquid nitrogen, and 2) all washing steps of the IgG-Sepharose and streptavidin-Sepharose Poly-Prep columns were performed using a peristaltic pump at a flow rate of 1 ml/min at 4°C.

### Mass spectrometry (MS) analysis of proteins co-purified with ARP6-TAP fusion protein

On-bead digestion of the TAP-purified proteins was performed as previously reported (Comstra et al., 2017; Cutler et al., 2017). Residual wash buffer was removed and 200 µl of 50 mM NH_4_HCO_3_ was added to each sample. Samples were reduced with 1 mM dithiothreitol for 30 minutes and alkylated with 5mM iodoacetamide in the dark for an additional 30 minutes. Both steps were performed at room temperature. Digestion was started with the addition of 1 µg of lysyl endopeptidase (Wako) for 2 hours and further digested overnight with 1:50 (w/w) trypsin (Promega) at room temperature. Resulting peptides were acidified with 25ul of 10% formic acid (FA) and 1% triflouroacetic acid (TFA), desalted with a Sep-Pak C18 column (Waters), and dried under vacuum.

Liquid chromatography coupled to tandem mass spectrometry (LC-MS/MS) on an Orbitrap Fusion mass spectrometer (ThermoFisher Scientific, San Jose, CA) was performed at the Emory Integrated Proteomics Core (EIPC) (Comstra et al., 2017; Cutler et al., 2017). The dried samples were resuspended in 10 µL of loading buffer (0.1% formic acid, 0.03% trifluoroacetic acid, 1% acetonitrile). Peptide mixtures (2 µL) were loaded onto a 25 cm x 75 µm internal diameter fused silica column (New Objective, Woburn, MA) self-packed with 1.9 µm C18 resin (Dr. Maisch, Germany). Separation was carried out over a 140-minute gradient by a Dionex Ultimate 3000 RSLCnano at a flow rate of 300 nL/min. The gradient ranged from 3% to 99% buffer B (buffer A: 0.1% formic acid in water, buffer B: 0.1 % formic in ACN). The spectrometer was operated in top speed mode with 3 second cycles. Full MS scans were collected in profile mode at 120,000 resolution at m/z 200 with an automatic gain control (AGC) of 200,000 and a maximum ion injection time of 50 ms. The full mass range was set from 400-1600 m/z. Tandem MS/MS scans were collected in the ion trap after higher-energy collisional dissociation (HCD). The precursor ions were isolated with a 0.7 m/z window and fragmented with 32% collision energy. The product ions were collected with the AGC set for 10,000 and the maximum injection time set to 35 ms. Previously sequenced precursor ions within +/- 10 ppm were excluded from sequencing for 20s using the dynamic exclusion parameters and only precursors with charge states between 2+ and 6+ were allowed.

All raw data files were processed using the Proteome Discoverer 2.1 data analysis suite (Thermo Scientific, San Jose, CA). The database was downloaded from Uniprot and consists of 33,388 *Arabidopsis thaliana* target sequences. An additional sequence was added for the TAP-tagged bait protein. Peptide matches were restricted to fully tryptic cleavage and precursor mass tolerances of +/- 20 ppm and product mass tolerances of +/- 0.6 Daltons. Dynamic modifications were set for methionine oxidation (+15.99492 Da) and protein N-terminal acetylation (+42.03670). A maximum of 3 dynamic modifications were allowed per peptide and a static modification of +57.021465 Da was set for carbamidomethyl cysteine. The Percolator node within Proteome Discoverer was used to filter the peptide spectral match (PSM) false discovery rate to 1%.

### Chromatin Immunoprecipitation (ChIP) with H2A.Z and H4Ac antibodies

For each sample, 1.5 grams of 6-days-old seedlings (without roots) were used for ChIP-seq and ChIP-qPCR experiments. For ChIP-seq experiments, 1.5 µg of the affinity-purified H2A.Z antibody (Deal et al., 2007) and 5 µl of H4Ac antibody (Millipore cat.# 06-866) were used. The ChIP-seq experiments were performed in biological triplicates on WT, *arp6-1*, and *mbd9-1* seedling tissues as described previously (Gendrel et al., 2005). The ChIP-qPCR experiment was performed in duplicates on WT, *arp6-1, mbd9-1, mbd9-2*, and *mbd9-3* seedling tissues as described previously (Gendrel et al., 2005), with the following modifications: 1) after the centrifugation of the nuclei in extraction buffer 3, the pellets were resuspended in 100 µl of nuclei lysis buffer, and 2) after sonication using a Diagenode Bioruptor, 100 µl of the fragmented chromatin was diluted with 1 ml of the ChIP dilution buffer and the whole solution was used for incubation with 1.5 µg of the affinity-purified H2A.Z antibody. The ChIP and input DNA samples from the ChIP-qPCR experiment were analyzed by real-time qPCR using the *ACT2* (*At3g18780*) 3’ untranslated region sequence as the endogenous control and with primers that span the genomic regions of *FLC (At5g10140), ASK11* (*At4g34210)*, and *At4 (At5g03545)* genes. The sequences of these primers were previously described (Deal et al., 2007; Smith et al., 2010).

### ChIP-seq library preparation, sequencing, and data analysis

Libraries were prepared starting with 500 pg of ChIP or input DNA using the Swift Accel-NGS 2S Plus DNA library kit (Swift Biosciences) according to the manufacturer’s instructions. All libraries were pooled and sequenced using single-end 50 nt reads on an Illumina NextSeq 500 instrument. Reads were mapped to the *Arabidopsis thaliana* genome (TAIR10) using the Bowtie2 package (Langmead and Salzberg, 2012). Quality filtering and sorting of the mapped reads, as well as removal of the reads that mapped to the organellar genomes was done as previously described (Sijacic et al., 2018) using Samtools 0.1.19 (Li et al., 2009). The filtered and sorted BAM files were converted to bigwig format as previously described (Sijacic et al., 2018) using deepTools 2.0 software (Ramirez et al., 2016). For visualization, for a given antibody, BAM files of each genotype were scaled to the same number of reads. Three scaled, replicate BAM files of each genotype for H2A.Z were combined and converted to a single bigwig file for each genotype. The same was done for H4A.C except that two scaled, replicates were combined for each genotype. Average plots displaying ChIP-seq data in Figures 2 and 4 were generated using the SeqPlots app (Stempor and Ahringer, 2016), while the Scatter plots, Heatmaps and average plots in Figure 3 were generated using “*computeMatrix*” “*plotHeatmap*” and “*plotProfile*” functions in the deepTools package.

### ChIP-seq peak calling

Peak calling on ChIP-seq data was done by employing the “*Findpeaks*” function of the HOMER package (Heinz et al., 2010) using the input ChIP-seq files as reference and the “*-region*” option. Called peaks were processed using Bedtools (Quinlan and Hall, 2010) to identify peaks called in at least one other replicate. This was done by keeping any peaks that overlapped by at least 50% between biological replicates. The retained peaks were concatenated and then merged together if they overlapped by at least 50%.

### Identification of MBD9-dependent and MBD9-independent H2A.Z sites

The number of H2A.Z ChIP-seq reads within the reproducible H2A.Z-enriched peaks from WT was quantified in WT, *mbd9-1*, and *arp6-1* plants using HTSeq’s *htseq-count* script (Anders et al., 2015). Three replicates of counted reads for all three genotypes were then processed using DESeq2 (Love et al., 2014). MBD9-dependent and MBD9- independent peaks were determined from the comparison between wild type and *mbd9-1* counted peaks. MBD9-dependent H2A.Z sites were identified as peaks that had a log fold change of 0.6 or more and an adjusted p-value less than or equal to 0.05. MBD9- independent H2A.Z sites were identified as peaks with an absolute log fold change less than 0.25.

### Genomic distribution of ChIP-seq peaks

The PAVIS web tool (Huang et al., 2013) was used to determine the genomic distribution of H2A.Z ChIP-seq peaks. The “upstream” regions were defined as the 2,000 bp upstream of the annotated transcription start site, and “downstream” regions were defined as the 1,000 bp downstream of the transcription end site.

### Identification of the genes nearest to the H2A.Z ChIP-seq peaks

Genes nearest to the MBD9-dependent and MBD9-independent H2A.Z sites were identified using the “*TSS*” function of the PeakAnnotator 1.4 program (Salmon-Divon et al., 2010) as previously described (Sijacic et al., 2018).

### Gene ontology analysis

Gene ontology (GO) analysis was carried out on gene lists from Supplemental Dataset 2 using two different GO web tools: 1) the AgriGO GO Analysis Toolkit, with default parameters (Du et al., 2010; Tian et al., 2017), and 2) Gene Ontology enrichment analysis (Mi et al., 2017). GO terms that had a false discovery rate (FDR) of 0.05 or less were considered significant.

### Violin plots of FPKM values

Publicly available RNA-seq FPKM values for genes nearest to the MBD9-dependent and MBD9-independent H2A.Z sites were plotted using ggplot2 (Wickham, 2016). Unpaired t-tests were used to determine whether FPKM values were significantly different between the two sets of genes. P values less than 0.05 were considered statistically significant.

### Real-time qPCR

Real-time qPCR was performed on the Applied Biosystems StepOnePlus real-time qPCR system using SYBR Green as a detection reagent. The 2^-ΔΔCt^ method (Livak and Schmittgen, 2001) of relative quantification was used to calculate the fold enrichment. The results presented for ChIP-qPCR experiments are average relative quantities from two biological replicates ± SD.

### Size-Exclusion Chromatography (SEC)

SEC was performed on the HiPrep 16/60 Sephacryl S-400 HR column (GE Healthcare) equilibrated with SEC buffer (the same extraction buffer as described in Van Leene et al., 2015, without NP-40 detergent). A mixture of protein standards ranging from 669 to 44 kDa (GE Healthcare), resuspended in the SEC buffer, were run on the column to produce a calibration curve of molecular weights versus elution volumes. The slope equation of the calibration curve was then used to calculate the molecular weight of the peak ARP6 SEC fractions. Total protein extracts were isolated from 1 gram of the WT and *mbd9-1* seedling tissues (without roots) using the same extraction buffer that was used for the ARP6-TAP-tag protein complex purification (Van Leene et al., 2015). For each run, between 1.8 and 2.0 ml of the protein extract was loaded onto the column and 1-ml fractions were collected. For each sample, two biological replicates of the SEC experiments were performed and gave nearly identical results.

### Publicly available ChIP-seq and RNA-seq data

Raw data from ChIP-seq experiments performed on young WT seedlings using antibodies against H3K4me3 (GSM2544796, (Wollmann et al., 2017)), H3K36me3 (GSM2544797, (Wollmann et al., 2017)), H3K9me2 (GSM2366607, (Jegu et al., 2017)), H3K9Ac (GSM2388452, (Kim et al., 2016)), H3K18Ac (GSM2096925, (Chen et al., 2017)), and H3K27Ac (GSM2096920, (Chen et al., 2017)) were processed and analyzed with the same procedures as for our ChIP-seq data (see above) and used to generate the average plots presented in Figure 4. The FPKM values from two different RNA-seq experiments (GSM2752981 and GSM2367133, respectively, (Lin et al., 2018; Zhou et al., 2017)), were used to compare expression levels in WT of genes nearest to the MBD9- dependent and MBD9-independent H2A.Z sites.

## Accession numbers

All ChIP-seq data generated in this study have been deposited to the NCBI GEO database under accession number GSE117391.

## Author contributions

P.S. and R.D. designed the research project. P.S., D.H., and R.D. carried out all experiments. M.B. and P.S. carried out data analyses. P.S. wrote the paper with input from all other authors.

## Acknowledgements

We thank Nicholas Seyfried and Duc Duong at the Emory Integrated Proteomics Core facility for carrying out the identification of ARP6-associated proteins. We also thank Eric Hoffer and Christine Dunham in the Emory Department of Biochemistry for their assistance with SEC experiments, and Grant Singer for assistance with plant genotyping. We would like to thank Jelle Van Leene and Geert Persiau for their technical suggestions regarding the TAP experiments. We are also grateful to the Arabidopsis Biological Resource Center at Ohio State University for providing the seeds of all T-DNA mutant lines.

## REFERENCES

Adam, M., Robert, F., Larochelle, M. and Gaudreau, L. (2001). H2A.Z is required for global chromatin integrity and for recruitment of RNA polymerase II under specific conditions. Mol Cell Biol 21, 6270–6279.

Altaf, M., Auger, A., Monnet-Saksouk, J., Brodeur, J., Piquet, S., Cramet, M., Bouchard, N., Lacoste, N., Utley, R., Gaudreau, L. and Cote, J. (2010). NuA4-dependent acetylation of nucleosomal histones H4 and H2A directly stimulates incorporation of H2A.Z by the SWR1 complex. J Biol Chem 285, 15966–15977.

Anders, S. and Huber, W. (2010). Differential expression analysis for sequence count data. Genome Biol 11, R106.

Anders, S., Pyl, P. T. and Huber, W. (2015). HTSeq-a Python framework to work with high-throughput sequencing data. Bioinformatics 31, 166–169.

Aravind, L. and Iyer, L. M. (2012). The HARE-HTH and associated domains: novel modules in the coordination of epigenetic DNA and protein modifications. Cell Cycle 11, 119–131.

Auger, A., Galarneau, L., Altaf, M., Nourani, A., Doyon, Y., Utley, R., Cronier, D., Allard, S. and Cote, J. (2008). Eaf1 is the platform for NuA4 molecular assembly that evolutionarily links chromatin acetylation to ATP-dependent exchange of histone H2A variants. Mol Cell Biol 28, 2257–2270.

Babiarz, J., Halley, J. and Rine, J. (2006). Telomeric heterochromatin boundaries require NuA4-dependent acetylation of histone variant H2A.Z in Saccharomyces cerevisiae. Genes Dev 20, 700–710.

Berg, A., Meza, T. J., Mahic, M., Thorstensen, T., Kristiansen, K. and Aalen, R. B. (2003). Ten members of the Arabidopsis gene family encoding methyl-CpG-binding domain proteins are transcriptionally active and at least one, AtMBD11, is crucial for normal development. Nucleic Acids Res 31, 5291–5304.

Bieluszewski, T., Galganski, L., Sura, W., Bieluszewska, A., Abram, M., Ludwikow, A., Ziolkowski, P. and Sadowski, J. (2015). AtEAF1 is a potential platform protein for Arabidopsis NuA4 acetyltransferase complex. BMC Plant Biol 15, 75.

Cai, Y., Jin, J., Florens, L., Swanson, S., Kusch, T., Li, B., Workman, J., Washburn, M., Conaway, R. and Conaway, J. (2005). The mammalian YL1 protein is a shared subunit of the TRRAP/TIP60 histone acetyltransferase and SRCAP complexes. J Biol Chem 280, 13665–13670.

Cai, Y., Jin, J., Gottschalk, A., Yao, T., Conaway, J. and Conaway, R. (2006). Purification and assay of the human INO80 and SRCAP chromatin remodeling complexes. Methods 40, 312–317.

Cai, Y., Jin, J., Tomomori-Sato, C., Sato, S., Sorokina, I., Parmely, T., Conaway, R. and Conaway, J. (2003). Identification of new subunits of the multiprotein mammalian TRRAP/TIP60-containing histone acetyltransferase complex. J Biol Chem 278, 42733–42736.

Chen, C., Li, C., Wang, Y., Renaud, J., Tian, G., Kambhampati, S., Saatian, B., Nguyen, I., Hannoufa, A., Marsolais, F., Yuan, Z. C., Yu, K., Austin, R. S., Liu, J., Kohalmi, S. E., Wu, K., Huang, S. and Cui, Y. (2017). Cytosolic acetyl-CoA promotes histone acetylation predominantly at H3K27 in Arabidopsis. Nat Plants 3, 814–824.

Cheng, X., Auger, A., Altaf, M., Drouin, S., Paquet, E., Utley, R. T., Robert, F. and Cote, J. (2015). Eaf1 Links the NuA4 Histone Acetyltransferase Complex to Htz1 Incorporation and Regulation of Purine Biosynthesis. Eukaryot Cell 14, 535–544.

Choi, K., Kim, J., Hwang, H., Kim, S., Park, C., Kim, S. and Lee, I. (2011). The FRIGIDA complex activates transcription of FLC, a strong flowering repressor in Arabidopsis, by recruiting chromatin modification factors. Plant Cell 23, 289–303.

Choi, K., Kim, S., Kim, S., Kim, M., Hyun, Y., Lee, H., Choe, S., Kim, S., Michaels, S. and Lee, I. (2005). SUPPRESSOR OF FRIGIDA3 encodes a nuclear ACTIN-RELATED PROTEIN6 required for floral repression in Arabidopsis. Plant Cell 17, 2647–2660.

Choi, K., Park, C., Lee, J., Oh, M., Noh, B. and Lee, I. (2007). Arabidopsis homologs of components of the SWR1 complex regulate flowering and plant development. Development 134, 1931–1941.

Clarkson, M. J., Wells, J. R., Gibson, F., Saint, R. and Tremethick, D. J. (1999). Regions of variant histone His2AvD required for Drosophila development. Nature 399, 694–697.

Clough, S. J. and Bent, A. F. (1998). Floral dip: a simplified method for Agrobacterium-mediated transformation of Arabidopsis thaliana. Plant J 16, 735–743.

Coleman-Derr, D. and Zilberman, D. (2012). Deposition of histone variant H2A.Z within gene bodies regulates responsive genes. PLOS Genet 8, e1002988.

Comstra, H. S., McArthy, J., Rudin-Rush, S., Hartwig, C., Gokhale, A., Zlatic, S. A., Blackburn, J. B., Werner, E., Petris, M., D’Souza, P., Panuwet, P., Barr, D. B., Lupashin, V., Vrailas-Mortimer, A. and Faundez, V. (2017). The interactome of the copper transporter ATP7A belongs to a network of neurodevelopmental and neurodegeneration factors. Elife 6

Conerly, M. L., Teves, S. S., Diolaiti, D., Ulrich, M., Eisenman, R. N. and Henikoff, S. (2010). Changes in H2A.Z occupancy and DNA methylation during B-cell lymphomagenesis. Genome Res 20, 1383–1390.

Curtis, M. D. and Grossniklaus, U. (2003). A gateway cloning vector set for high-throughput functional analysis of genes in planta. Plant Physiol 133, 462–469.

Cutler, A. A., Dammer, E. B., Doung, D. M., Seyfried, N. T., Corbett, A. H. and Pavlath, G. K. (2017). Biochemical isolation of myonuclei employed to define changes to the myonuclear proteome that occur with aging. Aging Cell 16, 738–749.

Deal, R., Kandasamy, M., McKinney, E. and Meagher, R. (2005). The nuclear actin-related protein ARP6 is a pleiotropic developmental regulator required for the maintenance of FLOWERING LOCUS C expression and repression of flowering in Arabidopsis. Plant Cell 17, 2633–2646.

Deal, R., Topp, C., McKinney, E. and Meagher, R. (2007). Repression of flowering in Arabidopsis requires activation of FLOWERING LOCUS C expression by the histone variant H2A.Z. Plant Cell 19, 74–83.

Deal, R. B. and Henikoff, S. (2010). A simple method for gene expression and chromatin profiling of individual cell types within a tissue. Dev Cell 18, 1030–1040.

Doyon, Y., Selleck, W., Lane, W., Tan, S. and Cote, J. (2004). Structural and functional conservation of the NuA4 histone acetyltransferase complex from yeast to humans. Mol Cell Biol 24, 1884–1896.

Du, Q., Luu, P. L., Stirzaker, C. and Clark, S. J. (2015). Methyl-CpG-binding domain proteins: readers of the epigenome. Epigenomics 7, 1051–1073.

Du, Z., Zhou, X., Ling, Y., Zhang, Z. and Su, Z. (2010). agriGO: a GO analysis toolkit for the agricultural community. Nucleic Acids Res 38, W64–70.

Durant, M. and Pugh, B. F. (2007). NuA4-directed chromatin transactions throughout the Saccharomyces cerevisiae genome. Mol Cell Biol 27, 5327–5335.

Efroni, I., Han, S. K., Kim, H. J., Wu, M. F., Steiner, E., Birnbaum, K. D., Hong, J. C., Eshed, Y. and Wagner, D. (2013). Regulation of leaf maturation by chromatin-mediated modulation of cytokinin responses. Dev Cell 24, 438–445.

Faast, R., Thonglairoam, V., Schulz, T. C., Beall, J., Wells, J. R., Taylor, H., Matthaei, K., Rathjen, P. D., Tremethick, D. J. and Lyons, I. (2001). Histone variant H2A.Z is required for early mammalian development. Curr Biol 11, 1183–1187.

Gendrel, A. V., Lippman, Z., Martienssen, R. and Colot, V. (2005). Profiling histone modification patterns in plants using genomic tiling microarrays. Nat Methods 2, 213–218.

Gevry, N., Chan, H., Laflamme, L., Livingston, D. and Gaudreau, L. (2007). p21 transcription is regulated by differential localization of histone H2A.Z. Genes Dev 21, 1869–1881.

Gomez-Zambrano, A., Crevillen, P., Franco-Zorrilla, J. M., Lopez, J. A., Moreno-Romero, J., Roszak, P., Santos-Gonzalez, J., Jurado, S., Vazquez, J., Kohler, C., Solano, R., Pineiro, M. and Jarillo, J. A. (2018). Arabidopsis SWC4 Binds DNA and Recruits the SWR1 Complex to Modulate Histone H2A.Z Deposition at Key Regulatory Genes. Mol Plant 11, 815–832.

Hale, C. J., Potok, M. E., Lopez, J., Do, T., Liu, A., Gallego-Bartolome, J., Michaels, S. D. and Jacobsen, S. E. (2016). Identification of Multiple Proteins Coupling Transcriptional Gene Silencing to Genome Stability in Arabidopsis thaliana. PLOS Genet 12, e1006092.

Heinz, S., Benner, C., Spann, N., Bertolino, E., Lin, Y. C., Laslo, P., Cheng, J. X., Murre, C., Singh, H. and Glass, C. K. (2010). Simple combinations of lineage-determining transcription factors prime cis-regulatory elements required for macrophage and B cell identities. Mol Cell 38, 576–589.

Hota, S. and Bruneau, B. (2016). ATP-dependent chromatin remodeling during mammalian development. Development 143, 2882–2897.

Huang, W., Loganantharaj, R., Schroeder, B., Fargo, D. and Li, L. (2013). PAVIS: a tool for Peak Annotation and Visualization. Bioinformatics 29, 3097–3099.

Jackson, J. D. and Gorovsky, M. A. (2000). Histone H2A.Z has a conserved function that is distinct from that of the major H2A sequence variants. Nucleic Acids Res 28, 3811–3816.

Jarillo, J. and Pineiro, M. (2015). H2A.Z mediates different aspects of chromatin function and modulates flowering responses in Arabidopsis. Plant J 83, 96–109.

Jegu, T., Veluchamy, A., Ramirez-Prado, J. S., Rizzi-Paillet, C., Perez, M., Lhomme, A., Latrasse, D., Coleno, E., Vicaire, S., Legras, S., Jost, B., Rougee, M., Barneche, F., Bergounioux, C., Crespi, M., Mahfouz, M. M., Hirt, H., Raynaud, C. and Benhamed, M. (2017). The Arabidopsis SWI/SNF protein BAF60 mediates seedling growth control by modulating DNA accessibility. Genome Biol 18, 114.

Keogh, M., Mennella, T., Sawa, C., Berthelet, S., Krogan, N., Wolek, A., Podolny, V., Carpenter, L., Greenblatt, J., Baetz, K. and Buratowski, S. (2006). The Saccharomyces cerevisiae histone H2A variant Htz1 is acetylated by NuA4. Genes Dev 20, 660–665.

Kim, Y. J., Wang, R., Gao, L., Li, D., Xu, C., Mang, H., Jeon, J., Chen, X., Zhong, X., Kwak, J. M., Mo, B., Xiao, L. and Chen, X. (2016). POWERDRESS and HDA9 interact and promote histone H3 deacetylation at specific genomic sites in Arabidopsis. Proc Natl Acad Sci U S A 113, 14858–14863.

Kimura, A. and Horikoshi, M. (1998). Tip60 acetylates six lysines of a specific class in core histones in vitro. Genes Cells 3, 789–800.

Kobor, M., Venkatasubrahmanyam, S., Meneghini, M., Gin, J., Jennings, J., Link, A., Madhani, H. and Rine, J. (2004). A protein complex containing the conserved Swi2/Snf2-related ATPase Swr1p deposits histone variant H2A.Z into euchromatin. PLOS Biol 2, 0588–0599.

Krogan, N., Keogh, M., Datta, N., Sawa, C., Ryan, O., Ding, H., Haw, R., Pootoolal, J., Tong, A., Canadien, V., Richards, D., Wu, X., Emili, A., Hughes, T., Buratowski, S. and Greenblatt, J. (2003). A Snf2 family ATPase complex required for recruitment of the histone H2A variant Htz1. Mol Cell 12, 1565–1576.

Kusch, T., Florens, L., Macdonald, W., Swanson, S., Glaser, R., Yates, J., Abmayr, S., Washburn, M. and Workman, J. (2004). Acetylation by Tip60 is required for selective histone variant exchange at DNA lesions. Science 306, 2084–2087.

Ladurner, A. G., Inouye, C., Jain, R. and Tjian, R. (2003). Bromodomains mediate an acetyl-histone encoded antisilencing function at heterochromatin boundaries. Mol Cell 11, 365–376.

Langmead, B. and Salzberg, S. L. (2012). Fast gapped-read alignment with Bowtie 2. Nat Methods 9, 357–359.

Lazaro, A., Gomez-Zambrano, A., Lopez-Gonzalez, L., Pineiro, M. and Jarillo, J. (2008). Mutations in the Arabidopsis SWC6 gene, encoding a component of the SWR1 chromatin remodelling complex, accelerate flowering time and alter leaf and flower development. J Exp Bot 59, 653–666.

Li, H., Handsaker, B., Wysoker, A., Fennell, T., Ruan, J., Homer, N., Marth, G., Abecasis, G. and Durbin, R. (2009). The Sequence Alignment/Map format and SAMtools. Bioinformatics 25, 2078–2079.

Li, H., Ilin, S., Wang, W., Duncan, E. M., Wysocka, J., Allis, C. D. and Patel, D. J. (2006). Molecular basis for site-specific read-out of histone H3K4me3 by the BPTF PHD finger of NURF. Nature 442, 91–95.

Lin, J. S., Kuo, C. C., Yang, I. C., Tsai, W. A., Shen, Y. H., Lin, C. C., Liang, Y. C., Li, Y. C., Kuo, Y. W., King, Y. C., Lai, H. M. and Jeng, S. T. (2018). MicroRNA160 Modulates Plant Development and Heat Shock Protein Gene Expression to Mediate Heat Tolerance in Arabidopsis. Front Plant Sci 9, 68.

Liu, X., Li, B. and Gorovsky Ma (1996). Essential and nonessential histone H2A variants in Tetrahymena thermophila. Mol Cell Biol 16, 4305–4311.

Livak, K. J. and Schmittgen, T. D. (2001). Analysis of relative gene expression data using real-time quantitative PCR and the 2(-Delta Delta C(T)) Method. Methods 25, 402–408.

Love, M. I., Huber, W. and Anders, S. (2014). Moderated estimation of fold change and dispersion for RNA-seq data with DESeq2. Genome Biol 15, 550.

Lu, P., Levesque, N. and Kobor, M. (2009). NuA4 and SWR1-C: two chromatin-modifying complexes with overlapping functions and components. Biochem Cell Biol 87, 799–815.

Luk, E., Ranjan, A., Fitzgerald, P. C., Mizuguchi, G., Huang, Y., Wei, D. and Wu, C. (2010). Stepwise histone replacement by SWR1 requires dual activation with histone H2A.Z and canonical nucleosome. Cell 143, 725–736.

March-Diaz, R., Garcia-Dominguez, M., Florencio, F. and Reyes, J. (2007). SEF, a new protein required for flowering repression in Arabidopsis, interacts with PIE1 and ARP6. Plant Physiol 143, 893–901.

March-Diaz, R., Garcia-Dominguez, M., Lozano-Juste, J., Leon, J., Florencio, F. and Reyes, J. (2008). Histone H2A.Z and homologues of components of the SWR1 complex are required to control immunity in Arabidopsis. Plant J 53, 475–487.

March-Diaz, R. and Reyes, J. (2009). The beauty of being a variant: H2A.Z and the SWR1 complex in plants. Mol Plant 2, 565–577.

Marques, M., Laflamme, L., Gervais, A. L. and Gaudreau, L. (2010). Reconciling the positive and negative roles of histone H2A.Z in gene transcription. Epigenetics 5, 267–272.

Martin-Trillo, M., Lazaro, A., Poethig, R., Gomez-Mena, C., Pineiro, M., Martinez-Zapater, J. and Jarillo, J. (2006). EARLY IN SHORT DAYS 1 (ESD1) encodes ACTIN-RELATED PROTEIN 6 (AtARP6), a putative component of chromatin remodelling complexes that positively regulates FLC accumulation in Arabidopsis. Development 133, 1241–1252.

Martinato, F., Cesaroni, M., Amati, B. and Guccione, E. (2008). Analysis of Myc-induced histone modifications on target chromatin. PLOS One 3, e3650.

Meneghini, M. D., Wu, M. and Madhani, H. D. (2003). Conserved histone variant H2A.Z protects euchromatin from the ectopic spread of silent heterochromatin. Cell 112, 725–736.

Mi, H., Huang, X., Muruganujan, A., Tang, H., Mills, C., Kang, D. and Thomas, P. D. (2017). PANTHER version 11: expanded annotation data from Gene Ontology and Reactome pathways, and data analysis tool enhancements. Nucleic Acids Res 45, D183–d189.

Millar, C., Xu, F., Zhang, K. and Grunstein, M. (2006). Acetylation of H2AZ Lys 14 is associated with genome-wide gene activity in yeast. Genes Dev 20, 711–722.

Mizuguchi, G., Shen, X., Landry, J., Wu, W., Sen, S. and Wu, C. (2004). ATP-driven exchange of histone H2AZ variant catalyzed by SWR1 chromatin remodeling complex. Science 303, 343–348.

Murashige, T. and Skoog, F. (1962). A revised medium for rapid growth and bio assays with tobacco tissue cultures. Physiologia Plantarum 15, 473–497.

Nguyen, V. Q., Ranjan, A., Stengel, F., Wei, D., Aebersold, R., Wu, C. and Leschziner, A. E. (2013). Molecular architecture of the ATP-dependent chromatin-remodeling complex SWR1. Cell 154, 1220–1231.

Noh, Y. and Amasino, R. (2003). PIE1, an ISWI family gene, is required for FLC activation and floral repression in Arabidopsis. Plant Cell 15, 1671–1682.

Peng, M., Cui, Y., Bi, Y. and Rothstein, S. (2006). AtMBD9: a protein with a methyl-CpG-binding domain regulates flowering time and shoot branching in Arabidopsis. Plant J 46, 282–296.

Quinlan, A. R. and Hall, I. M. (2010). BEDTools: a flexible suite of utilities for comparing genomic features. Bioinformatics 26, 841–842.

Raisner, R. M. and Madhani, H. D. (2006). Patterning chromatin: form and function for H2A.Z variant nucleosomes. Curr Opin Genet Dev 16, 119–124.

Ramirez, F., Ryan, D. P., Gruning, B., Bhardwaj, V., Kilpert, F., Richter, A. S., Heyne, S., Dundar, F. and Manke, T. (2016). deepTools2: a next generation web server for deep-sequencing data analysis. Nucleic Acids Res 44, W160–165.

Rosa, M., Von Harder, M., Cigliano, R. A., Schlogelhofer, P. and Mittelsten Scheid, O. (2013). The Arabidopsis SWR1 chromatin-remodeling complex is important for DNA repair, somatic recombination, and meiosis. Plant Cell 25, 1990–2001.

Ruhl, D., Jin, J., Cai, Y., Swanson, S., Florens, L., Washburn, M., Conaway, R., Conaway, J. and Chrivia, J. (2006). Purification of a human SRCAP complex that remodels chromatin by incorporating the histone variant H2A.Z into nucleosomes. Biochemistry 45, 5671–5677.

Salmon-Divon, M., Dvinge, H., Tammoja, K. and Bertone, P. (2010). PeakAnalyzer: genome-wide annotation of chromatin binding and modification loci. BMC Bioinformatics 11, 415.

Scebba, F., Bernacchia, G., De Bastiani, M., Evangelista, M., Cantoni, R. M., Cella, R., Locci, M. T. and Pitto, L. (2003). Arabidopsis MBD proteins show different binding specificities and nuclear localization. Plant Mol Biol 53, 715–731.

Schmid, M., Davison, T., Henz, S., Pape, U., Demar, M., Vingron, M., Scholkopf, B., Weigel, D. and Lohmann, J. (2005). A gene expression map of Arabidopsis thaliana development. Nat Genet 37, 501–506.

Sijacic, P., Bajic, M., McKinney, E. C., Meagher, R. B. and Deal, R. B. (2018). Changes in chromatin accessibility between Arabidopsis stem cells and mesophyll cells illuminate cell type-specific transcription factor networks. Plant J 94, 215–231.

Smith, A. P., Jain, A., Deal, R. B., Nagarajan, V. K., Poling, M. D., Raghothama, K. G. and Meagher, R. B. (2010). Histone H2A.Z regulates the expression of several classes of phosphate starvation response genes but not as a transcriptional activator. Plant Physiol 152, 217–225.

Stempor, P. and Ahringer, J. (2016). SeqPlots - Interactive software for exploratory data analyses, pattern discovery and visualization in genomics. Wellcome Open Res 1, 14.

Strohner, R., Nemeth, A., Jansa, P., Hofmann-Rohrer, U., Santoro, R., Langst, G. and Grummt, I. (2001). NoRC-a novel member of mammalian ISWI-containing chromatin remodeling machines. Embo j 20, 4892–4900.

Surface, L. E., Fields, P. A., Subramanian, V., Behmer, R., Udeshi, N., Peach, S. E., Carr, S. A., Jaffe, J. D. and Boyer, L. A. (2016). H2A.Z.1 Monoubiquitylation Antagonizes BRD2 to Maintain Poised Chromatin in ESCs. Cell Rep 14, 1142–1155.

Tian, T., Liu, Y., Yan, H., You, Q., Yi, X., Du, Z., Xu, W. and Su, Z. (2017). agriGO v2.0: a GO analysis toolkit for the agricultural community, 2017 update. Nucleic Acids Res 45, W122–w129.

van Daal, A. and Elgin, S. C. (1992). A histone variant, H2AvD, is essential in Drosophila melanogaster. Mol Biol Cell 3, 593–602.

Van Leene, J., Eeckhout, D., Cannoot, B., De Winne, N., Persiau, G., Van De Slijke, E., Vercruysse, L., Dedecker, M., Verkest, A., Vandepoele, K., Martens, L., Witters, E., Gevaert, K. and De Jaeger, G. (2015). An improved toolbox to unravel the plant cellular machinery by tandem affinity purification of Arabidopsis protein complexes. Nat Protoc 10, 169–187.

Vercruyssen, L., Verkest, A., Gonzalez, N., Heyndrickx, K. S., Eeckhout, D., Han, S. K., Jegu, T., Archacki, R., Van Leene, J., Andriankaja, M., De Bodt, S., Abeel, T., Coppens, F., Dhondt, S., De Milde, L., Vermeersch, M., Maleux, K., Gevaert, K., Jerzmanowski, A., Benhamed, M., Wagner, D., Vandepoele, K., De Jaeger, G. and Inze, D. (2014). ANGUSTIFOLIA3 binds to SWI/SNF chromatin remodeling complexes to regulate transcription during Arabidopsis leaf development. Plant Cell 26, 210–229.

Wei, W., Zhang, Y., Tao, J., Chen, H., Li, Q., Zhang, W., Ma, B., Lin, Q., Zhang, J. and Chen, S. (2015). The Alfin-like homeodomain finger protein AL5 suppresses multiple negative factors to confer abiotic stress tolerance in Arabidopsis. Plant J 81, 871–883.

Wollmann, H., Stroud, H., Yelagandula, R., Tarutani, Y., Jiang, D., Jing, L., Jamge, B., Takeuchi, H., Holec, S., Nie, X., Kakutani, T., Jacobsen, S. E. and Berger, F. (2017). The histone H3 variant H3.3 regulates gene body DNA methylation in Arabidopsis thaliana. Genome Biol 18, 94.

Wong, M., Cox, L. and Chrivia, J. (2007). The chromatin remodeling protein, SRCAP, is critical for deposition of the histone variant H2A.Z at promoters. J Biol Chem 282, 26132–26139.

Wu, M. F., Sang, Y., Bezhani, S., Yamaguchi, N., Han, S. K., Li, Z., Su, Y., Slewinski, T. L. and Wagner, D. (2012). SWI2/SNF2 chromatin remodeling ATPases overcome polycomb repression and control floral organ identity with the LEAFY and SEPALLATA3 transcription factors. Proc Natl Acad Sci U S A 109, 3576–3581.

Wu, W., Wu, C., Ladurner, A., Mizuguchi, G., Wei, D., Xiao, H., Luk, E., Ranjan, A. and Wu, C. (2009). N terminus of Swr1 binds to histone H2AZ and provides a platform for subunit assembly in the chromatin remodeling complex. J Biol Chem 284, 6200–6207.

Xiao, J., Jin, R., Yu, X., Shen, M., Wagner, J. D., Pai, A., Song, C., Zhuang, M., Klasfeld, S., He, C., Santos, A. M., Helliwell, C., Pruneda-Paz, J. L., Kay, S. A., Lin, X., Cui, S., Garcia, M. F., Clarenz, O., Goodrich, J., Zhang, X., Austin, R. S., Bonasio, R. and Wagner, D. (2017). Cis and trans determinants of epigenetic silencing by Polycomb repressive complex 2 in Arabidopsis. Nat Genet 49, 1546–1552.

Xu, Y., Ayrapetov, M. K., Xu, C., Gursoy-Yuzugullu, O., Hu, Y. and Price, B. D. (2012). Histone H2A.Z controls a critical chromatin remodeling step required for DNA double-strand break repair. Mol Cell 48, 723–733.

Yaish, M., Peng, M. and Rothstein, S. (2009). AtMBD9 modulates Arabidopsis development through the dual epigenetic pathways of DNA methylation and histone acetylation. Plant J 59, 123–135.

Yang, X. J. (2004). Lysine acetylation and the bromodomain: a new partnership for signaling. Bioessays 26, 1076–1087.

Zemach, A. and Grafi, G. (2007). Methyl-CpG-binding domain proteins in plants: interpreters of DNA methylation. Trends Plant Sci 12, 80–85.

Zemach, A., McDaniel, I. E., Silva, P. and Zilberman, D. (2010). Genome-wide evolutionary analysis of eukaryotic DNA methylation. Science 328, 916–919.

Zhang, H., Richardson, D. O., Roberts, D. N., Utley, R., Erdjument-Bromage, H., Tempst, P., Cote, J. and Cairns, B. R. (2004). The Yaf9 component of the SWR1 and NuA4 complexes is required for proper gene expression, histone H4 acetylation, and Htz1 replacement near telomeres. Mol Cell Biol 24, 9424–9436.

Zhou, B. O., Wang, S. S., Xu, L. X., Meng, F. L., Xuan, Y. J., Duan, Y. M., Wang, J. Y., Hu, H., Dong, X., Ding, J. and Zhou, J. Q. (2010). SWR1 complex poises heterochromatin boundaries for antisilencing activity propagation. Mol Cell Biol 30, 2391–2400.

Zhou, Y., Romero-Campero, F. J., Gomez-Zambrano, A., Turck, F. and Calonje, M. (2017). H2A monoubiquitination in Arabidopsis thaliana is generally independent of LHP1 and PRC2 activity. Genome Biol 18, 69.

Zilberman, D., Coleman-Derr, D., Ballinger, T. and Henikoff, S. (2008). Histone H2A.Z and DNA methylation are mutually antagonistic chromatin marks. Nature 456, 125–129.

Zlatanova, J. and Thakar, A. (2008). H2A.Z: view from the top. Structure 16, 166–179.

